# Strikingly different roles of SARS-CoV-2 fusion peptides uncovered by neutron scattering

**DOI:** 10.1101/2021.08.30.458099

**Authors:** Andreas Santamaria, Krishna C. Batchu, Olga Matsarskaia, Sylvain F. Prévost, Daniela Russo, Francesca Natali, Tilo Seydel, Ingo Hoffmann, Valerie Laux, Michael Haertlein, Tamim A. Darwish, Robert A. Russell, Giacomo Corucci, Giovanna Fragneto, Armando Maestro, Nathan R. Zaccai

## Abstract

Coronavirus disease-2019 (COVID-19), a lethal respiratory illness caused by the coronavirus SARS-CoV-2, emerged in the end of 2019 and has since spread aggressively across the globe. A thorough understanding of the molecular mechanisms of cellular infection by coronaviruses is therefore of utmost importance. A critical stage in infection is the fusion between viral and host membranes. Here, we present a detailed investigation of the role of the SARS-CoV-2 Spike protein, and the influence of calcium and cholesterol, in this fusion process. Structural information from specular neutron reflectometry and small angle neutron scattering, complemented by dynamics information from quasi-elastic and spin-echo neutron spectroscopy, revealed strikingly different functions encoded in the Spike fusion domain. Calcium drives the N-terminal of the Spike fusion domain to fully cross the host plasma membrane. Removing calcium however re-orients the protein to the lipid leaflet in contact with the virus, leading to significant changes in lipid fluidity and rigidity. In conjunction with other regions of the fusion domain which are also positioned to bridge and dehydrate viral and host membranes, the molecular events leading to cell entry by SARS-CoV-2 are proposed.

**SIGNIFICANCE:** We have recreated here important elements of the critical membrane fusion mechanism of the SARS-CoV-2 coronavirus by simplifying the system down to its core elements, amenable to experimental analysis by neutron scattering. Neutrons are well suited for the study of protein – membrane interactions under physiological conditions, since they allow structural and dynamics characterization at room temperature. Our results revealed strikingly different functions encoded in the viral Spike fusion domain and thereby provide a potential calcium-dependent cell entry mechanism for SARS-CoV-2. In particular, calcium drives the protein’s N-terminal to harpoon through the host membrane, while removing calcium re-orients the protein so that it is able to bridge and dehydrate lipid membranes, facilitating their fusion.

## INTRODUCTION

β-coronaviruses (CoVs) are single-stranded positive sense RNA viruses (1). A deadly outbreak of respiratory tract infections caused by the SARS-CoV-2 β-coronavirus emerged in late 2019 in China (2, 3) and was declared a worldwide pandemic in March 2020. Other β-coronaviruses, including severe acute respiratory syndrome coronavirus 1 (SARS-CoV-1) and Middle Eastern respiratory syndrome coronavirus (MERS-CoV) are also highly contagious pathogens. A thorough understanding of the molecular mechanisms of cellular infection by coronaviruses is therefore of utmost importance.

The main structural components of β-coronaviruses include a lipid envelope, the Spike, Membrane and Envelope proteins, as well as the Nucleoproteins, which form a complex with the viral RNA. The Spike protein has been directly implicated in SARS-CoV-2 infectivity (4-6) and is consequently a target for both vaccine and antiviral drug efforts (7, 8). In a mechanism also present in other RNA viruses, like HIV and the influenza virus (9), this class I viral fusion glycoprotein induces fusion between the viral and host cellular lipid membranes, thereby facilitating viral entry. The focus of this work is to provide a physical and thermodynamic description of the molecular events leading up to fusion between SARS-CoV-2 and host membranes. Moreover, cell entry of SARS-CoV-2 can either occur directly through fusion at the plasma membrane (10) (in the presence of calcium), or after endocytosis, at the endosomal membrane (6) (where free calcium concentrations are greatly reduced (11)). Therefore, this study also focused on the influence of calcium on the fusion of cholesterol-rich membranes, such as the plasma membrane.

The Spike protein’s N-terminal S1 subunit contains the receptor-binding domain for the angiotensin-converting enzyme 2 (ACE-II). After binding, a proteolysis-triggered conformational change in the C-terminal S2 subunit mediates fusion between the viral and host cell membranes (6, 12, 13). S2 continues to be embedded in the viral membrane, but its heptad repeat 1 (HR1) and 2 (HR2) domains associate to form a six-helix bundle fusion core (8). Proteolysis at the S2’ site (at residue 816) subsequently frees the Spike protein fusion domain to associate with the host cell and initiate membrane fusion.

The Spike protein membrane fusion domain is strongly conserved between β-coronaviruses (7). However, the exact “fusion peptide” at or near the newly generated N-terminus of S2’, has not yet been conclusively identified. The expected characteristics - short, hydrophobic, with a possible canonical fusion tripeptide (YFG or FXG) along with a central proline residue (14) suggest several putative fusion peptides: FP1 (SARS-CoV-2 816-837), FP2 (835-856), FP3 (854-874) and FP4 (885-909) (also named internal fusion peptide (15)).

In SARS-CoV-1 and MERS-CoV, mutagenesis of the highly conserved S2′-proximal FP1 in the context of full-length Spike protein demonstrated its importance in mediating membrane fusion (16). SARS-CoV-2 FP1 retains the conserved LLF motif, which was critical for SARS-CoV-1 membrane fusion (17, 18). FP2 is characterized by a pair of highly conserved cysteines, which if reduced in SARS-CoV-1, lead to the abrogation of their membrane-ordering effect. Importantly, SARS-CoV-1 and -2 peptides containing FP1 and FP2 are able to induce membrane ordering in a Ca^2+^-dependent fashion (15, 18, 19). FP3 has an FXG motif at its N-terminus, and near its center, two prolines are surrounded by hydrophobic residues. Adjacent to HR1, FP4 has a central proline and a C-terminal FXG motif. Present within the peptide, the conserved SARS-CoV-1 and -2 GAALQIPFAMQMAYRF sequence can induce hemifusion between small unilamellar vesicles (15), although the peptide does not induce membrane ordering (18).

In this work, several neutron scattering techniques were employed to study the structure and dynamics of the interaction of fusion peptides with model membrane systems (see **Tables S1** and **S2**, respectively). Neutrons are particularly well suited for the study of soft and biological matter since they allow measurements at room-temperature with a fraction of nanometer resolution and at energies corresponding to thermal fluctuations. They are non-destructive and highly penetrating, thus allowing work in physiological conditions. Furthermore, as neutrons interact very differently with hydrogen (^1^H) and deuterium (^2^H), it is possible through isotopic substitution, to observe hydrogen atoms and water molecules in biological samples, and therefore highlight structural and chemical differences in specific regions of interest.

Different *in vitro* models of increasing complexity were used to characterize plasma membrane (PM) models interacting with SARS-CoV-2 peptides. The goal was to recreate important elements of the viral fusion mechanism by simplifying the system down to its core elements. For example, even though solid-supported lipid bilayers are appropriate model membrane systems, a lipid monolayer already provides a simple and versatile model for peptide assembly and insertion into a lipid membrane. The air/buffer interface mimics the opposite membrane leaflet, while by using a Langmuir trough, lateral membrane surface pressure can be controlled by restricting the monolayer lipids to a specific surface area, thereby recreating the pressure experienced within a lipid bilayer. Importantly, synthetic model membranes, as well as biomimetic plasma membrane (PM) with physiological phospholipid and cholesterol composition were studied. Depleting membrane cholesterol inhibits viral membrane fusion by SARS-CoV-2 and other coronaviruses (20). No chemical moieties were added to the lipids (or to the peptides), as an important advantage of neutrons in the study of lipid membranes is that the experiments are label-free. Similarly, to avoid problems such as artifactual interactions with the biomimetic PM lipids, the peptides were not modified by introducing additional residues, such as lysine-rich hydrophilic tags.

Through neutron scattering and biophysical experiments, physical and thermodynamic aspects of the membrane fusion mechanism of SARS-CoV-2 were investigated. Structural information from Specular Neutron Reflectometry (SNR) and Small Angle Neutron Scattering (SANS) were complemented by dynamics information from Quasi-Elastic (QENS) and Spin-Echo (NSE) neutron scattering accessing membrane fluidity and rigidity. The data highlight strikingly different roles for the different regions of the SARS-CoV-2 Spike protein fusion domain.

## RESULTS

### A biomimetic plasma membrane

The eukaryotic PM is characterized by a very high cholesterol content (∼50% mol%) (21) (**Table S2**). At room temperature, the surface pressure (Π) - Area (A) isotherm did not show any phase transition, suggesting that the biomimetic PM monolayer is in the liquid expanded (LE) phase (**Figure S1 A**). At a surface pressure of Π ≈ 25 mN m^−1^, which is the typical lateral pressure of a lipid membrane bilayer (21, 22), the in-plane structure of the PM monolayer obtained by BAM shows the coexistence of a densely packed liquid-ordered (Lo) phase (cholesterol-rich, brighter regions) and liquid-disordered (Ld) domains with a high lateral diffusion of lipids (**Figure 1 A, and S1 B**). The observed phase separation in lipid-only membranes was previously reported by several laboratories (as reviewed by Marsh (23). The vertical structure of PM, perpendicular to the plane of the interface, was determined by SNR analysis of hydrogenous and deuterated PM lipid monolayers in two different D2O buffers (**Figure S1 C, D** and **Table S3**). PM monolayers were effectively modelled as two layers: the 16 ± 1Å long aliphatic lipid tails, which are exposed to air, and the 8 ± 1Å lipid head-groups, in contact with the buffer. The mean area per phospholipid molecule (APM) obtained was 60 ± 4Å^2^, which is similar to the value determined from the pressure-area isotherm (see Supplementary Information, **SI Methods**) and is also consistent with a compact lipid bilayer (24, 25). From the neutron scattering length density (SLD) distribution (see **SI Methods**), the volume fraction of the different PM components could be determined (**Figure S1 E**). The cholesterol molecules are confined to the aliphatic tails layer, with no solvent present. A solvent penetration of 63% occurs in the head-groups layer.

**Figure 1.**
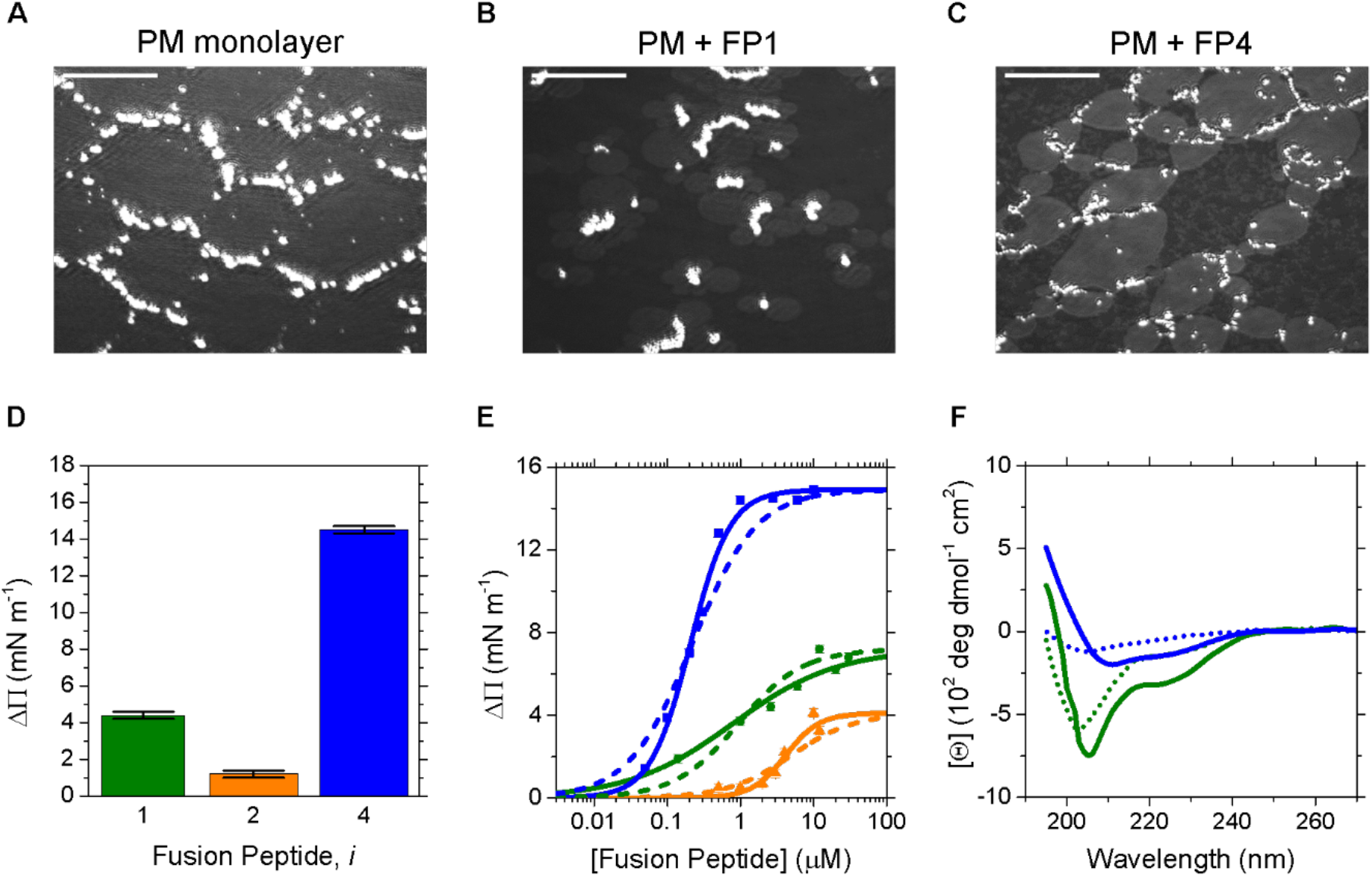
Interaction of Fusion Peptides with biomimetic lipid monolayers. **A, B, C** BAM images of the PM monolayer before (at a surface pressure of 23± 1mN m^-1^) and after injection of either FP1 or of FP4. Scale bars are 50μm. **D, E** Tensiometry binding analysis for FP binding to the PM determined from lateral pressure changes in the lipid monolayer. **D** Uppermost surface pressure increment due to the interaction of FP1, FP2, FP3 and FP4 with a PM monolayer, at 3μM peptide concentration and Π_0_ = 23± 1mN m^-1^. The increment in the pressure, ΔΠ, is proportional to the amount of peptide partitioning with the monolayer at the air/water interface, either through binding and/or through insertion. **E** Equilibrium analyses and resultant K_Ds_ for FP1 (green), FP2 (orange) and FP4 (green) binding to PM. Lines are fits to the data obtained through the Hill-Langmuir equation, as described in **SI Methods, Equation S1**. The dotted lines indicate the fits of the experimental data with the Hill coefficient set to 1. **F** CD molar ellipticity profiles of FP1 (green) and FP4 (green), in the absence (dotted line) and presence of liposomes (continuous line). The decrease in signal in the presence of lipids suggests a related increase in protein secondary structure.

Solid-supported lipid bilayers subsequently analysed by SNR revealed a similar membrane organization, with the cholesterol integrated into the acyl layer. The average area per phospholipid molecule obtained (from 53 to 59Å^2^) and the total thickness of 47 ± 2Å agrees with similar systems studied (26). The coverage was not less than 90% confirming full lipid coverage (**Table S6**).

### Interaction of FP1, FP2, FP3 and FP4 with the PM

In the Langmuir trough, the injection of either FP1, FP2, FP3 or FP4 into the bulk buffer underneath the PM monolayer gives rise to an immediate and rapid increase in surface pressure, followed by a slower increase until a plateau is reached typically several hours later (**Figure S2 A)**. The obtained pressure increment, ΔΠ, in the presence of 3 µM peptide in the bulk buffer, is plotted in **Figure 1 D**. The binding affinity of the different FPs, in the absence of calcium, was therefore determined from the FP concentration and the resulting change in surface pressure (**Figure 1 E, SI Methods**). FP3 was not fully characterized due to its weak affinity. Uncharged FP2 had a dissociation constant KD 14 ± 8µM (Hill coefficient of 1.9 ± 0.5). The K_D_ of FP1 was 0.9 ± 0.1μM (Hill coefficient of 0.61 ± 0.06). A Hill coefficient lower than 1 indicates negative cooperative binding. It may be due to FP1’s overall negative charge at physiological pH, which is not counterbalanced on binding the non-negligible amounts of negatively charged PS lipids in the PM outer leaflet (27). FP4 had the highest affinity to the PM. Its K_D_ was 80nM ± 10nM. The Hill coefficient of 1.6 ± 0.1 is significantly higher than 1, indicating a positively cooperative binding, possibly due to FP4’s positive charge at acidic and physiological pH being compensated by binding negatively charged PM. The resulting binding free energy to PM monolayer of FP1, FP2 and FP4 were 8.2 kcal/mol, 6.6 kcal/mol, 9.7 kcal/mol respectively.

In the presence of cholesterol-rich liposomes, the decrease in CD signal between 220nm and 235nm indicates the presence of increased secondary structure for FP1 and FP4 peptides (**Figure 1F**). These observations were previously noted for FP1 with liposomes (19) and for sections of FP1 and FP4 in the lipid mimetic trifluoroethanol (17, 28).

The Lo/Ld coexistence in the PM liquid expanded phase was clearly perturbed by the presence of either FP1 or FP4 (**Figure 1 B, C**). The former yields a change in the in-plane morphology where bright spots, potentially representing cholesterol clusters, are now isolated but surrounded by clearly visible domains. In the case of FP4, the resultant bright spots are still interconnected and surrounded by more homogeneous regions.

### FP1 buries deep into the PM, while FP2 and FP4 bind lipid headgroups

Due to the weak effect of FP3 on the PM monolayer, SNR experiments focused on FP1, FP2 and FP4. **Figure S3** shows the vertical reflectivity profiles measured at 4 different isotopic contrasts after the binding of the peptides to the PM monolayer. Data modeling was performed by simultaneously fitting all contrasts to obtain a single set of structural parameters that allow us to determine the percentage of the volume fraction of peptide partitioning into the lipids (see **Figure 2 A, B, C Table S5**). Two-layers models, which included the partition of the FPs into the aliphatic lipid tails and lipid polar headgroups, were adequate to describe the experimental data (see **Figure S3)**. Including in the model a third layer for the peptides yielded a worse fit of the data and suggests that the interaction of FP1, FP2 and FP4 with the membrane is not due to a physisorption of the peptides, but rather to their insertion directly into the lipid monolayer.

**Figure 2.**
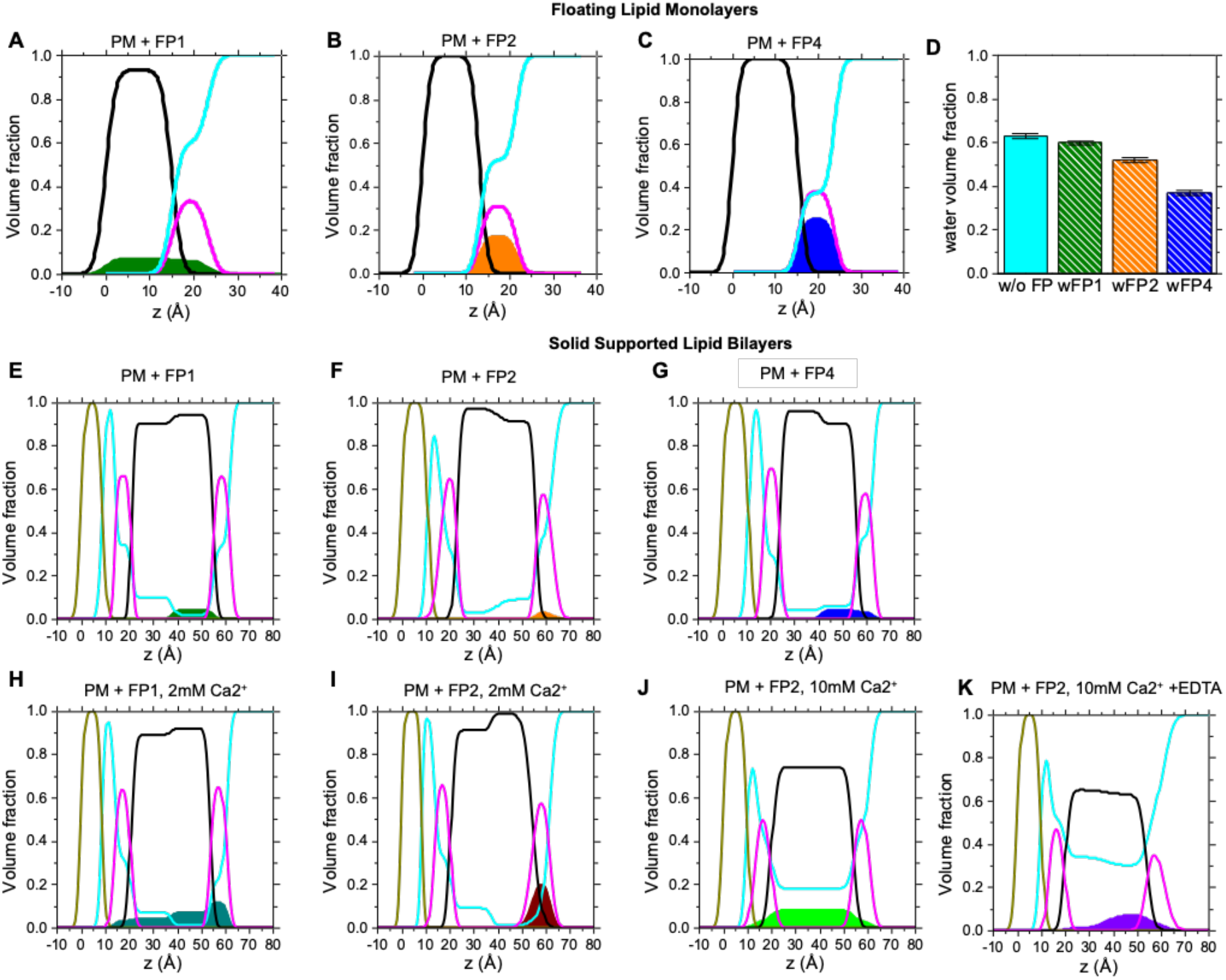
**A, B, C** Neutron reflectivity data of PM monolayers with FP1, FP2 and FP4 in the absence of Ca^2+^. Volume fraction profiles normal to the interface of PM monolayers (derived from data plotted in **Figure S3**, as described in **SI Methods**) highlight the distribution of tails (black), heads (magenta), water (cyan) and **A** FP1 (green), **B** FP2 (orange) and **C** FP4 (blue). **D** Bar diagram plot summarizing the volume fraction occupied by the solvent in the lipid headgroup layer of the PM monolayers with FP1, FP2 and FP4. **E, F, G** Neutron reflectivity data of PM bilayers with FP1, FP2 and FP4 in the absence of Ca^2+^. Volume fraction profiles normal to the interface of solid supported lipid bilayers (derived from data plotted in **Figure S4**, as described in **SI Methods**) highlight the distribution of Si/SO_2_ (grey), tails (black), heads (magenta), water (cyan) and **E** FP1 (green), **F** FP2 (orange), **G** FP4 (blue). **H, I, J** Neutron reflectivity data of PM bilayers with FP1 and FP2 in the presence of Ca^2+.^ (derived from data plotted in **Figure S5**). Volume fraction profiles normal to the interface of solid supported lipid bilayers are plotted with **H** FP1 with 2mM Ca^2+.^ (green) and **I** FP2 with 2mM Ca^2+.^ (maroon). **J, K** Neutron reflectivity data of PM bilayer with FP1 in the presence and then after removal of Ca^2+^. Volume fraction profiles normal to the interface of solid supported lipid bilayers are plotted with **J** FP4 with 10mM Ca^2+.^ (light green) and **K** FP4 after over-night incubation with EDTA (purple). Data from FIGARO at ILL.

The bound FP1 is found distributed across the entire PM monolayer, as it is present in both the lipid headgroups (6±1%) and aliphatic tails (7±1%) (**Figure 2 A**). In contrast, FP2 and FP4 had a negligible presence in the acyl region (**Figure 2 B, C**). FP2 and FP4 interacted more strongly within the lipid head region (17% for FP2 and 25% for FP4), where significant decreases in hydration were moreover observed (**Figure 2 D**).

Due to the manner in which FP1, FP2 and FP4 partitioned into the lipid monolayer, solid-supported bilayers, composed of synthetic lipids enriched in cholesterol, were also investigated (**Figure 2 E, F, G**). Bilayer integrity is conserved after FP binding, according to SNR (see **Figures S4 S5** and **Table S6**). As previously observed in the interaction of the Spike extra cellular domain with lipid bilayers (29), surface coverage was reduced, preventing direct comparison of lipid head group solvation. Again, the best-fit model did not require an additional peptide-rich layer. The changes observed in the tail region bilayer after the addition of FP1 are compatible with the partition of the peptide into the outer leaflet acyl region (4%). FP2 and FP4 are still positioned within the outer head groups (3% in both cases) although for FP4, its presence is also detected in the outer lipid acyl region (4%). The inner leaflet showed neither FP1, FP2 nor FP4 insertion.

### Calcium induces formation of a transmembrane FP1 across cholesterol-enriched lipid bilayers

The presence of calcium increases the membrane ordering effect of SARS-CoV-2 FP1-2 (18, 19). To this end, we studied the binding of FP1 and of FP2 in presence of Ca^2+^ and we found that, although FP2 still bound within the lipid head group region, the partition of FP1 into the membrane bilayer drastically changed (**Figure 2 H, I, J** and **S5 Table S6**).

In 2mM Ca^2+^, bound FP1 peptide was no longer limited to the outer leaflet but distributed across the whole lipid bilayer with 7% and 4% peptide in the outer and inner lipid tails region, respectively, and 12% and 3% in both the inner and outer headgroup layer. At a higher Ca^2+^ concentration (10 mM), an increase in the solvent fraction in both leaflets, from 34 to 43% solvent in the head groups region but also from 8% to 18% in the tail regions was also observed. The majority of FP1 was distributed in the tail region (8% versus 3% in both inner and outer head-group regions). Subsequently, adding the chelating agent EDTA to the bilayer to remove Ca^2+^ clearly reduced the amount of FP1 partitioned into the bilayer’s inner leaflet, whilst only slightly affecting the amount of FP1 partitioned in the outer tail region (7%) (**Figure 2 K**). The FP1 proportion in the outer headgroup region slightly increased from 3 to 4%. These data suggest that FP1 forms a calcium-dependent transmembrane peptide across the bilayer. When calcium is absent or removed by chelation, FP1 would only insert into the bilayer’s outer leaflet.

These SNR data also support the hypothesis that FP1 induces lipid reorganization. Upon removal of the calcium, the best fit bilayer model results in a significant increase in solvent percentage in the lipid tail region (from 17% to 30%).

In the case of FP2, in the presence of 2mM Ca^2+^, whilst no FP2 is observed in either the tail region or in the inner head group layer, there is a significant increase in peptide bound in the outer head-group layer, from 3% to 24% (**Figure 2 F, I**), which is tentatively linked to an increase in affinity to the negatively charged PM (27).

### FP1 disrupts liposomes, while FP4 induces liposome clustering

The effect on closed curved lipid bilayers of the different binding behavior between FP1, which inserts into the lipid tail region, and FP2 and FP4, which lie at the surface of the lipid bilayer, were investigated by SANS. In order to address the role of cholesterol, PM, ERGIC membrane (20% cholesterol) and “O” membrane (lacking cholesterol) were studied, at a constant concentration (∼1mg/ml), and across a wide range of peptide:lipid molar ratios (1:2000 to 1:6). The viral membrane would have a similar composition as the ERGIC membrane (30).

In the absence of any peptide, SUVs were stable over several weeks with a typical translucent appearance of the solution. SANS profiles were characteristic of SUVs with a form factor oscillation at high q, corresponding to a lamellar thickness of 3.4 nm for different cholesterol contents and FPs, followed by a q^-2^ power law indicating a flat SUV interface. (**Figure 3** and **S6**) The low-q onset of a plateau provided the overall size of the SUVs (120 nm in the absence of FPs is compatible with lipid extrusion through a 100 nm pore-size membrane).

**Figure 3.**
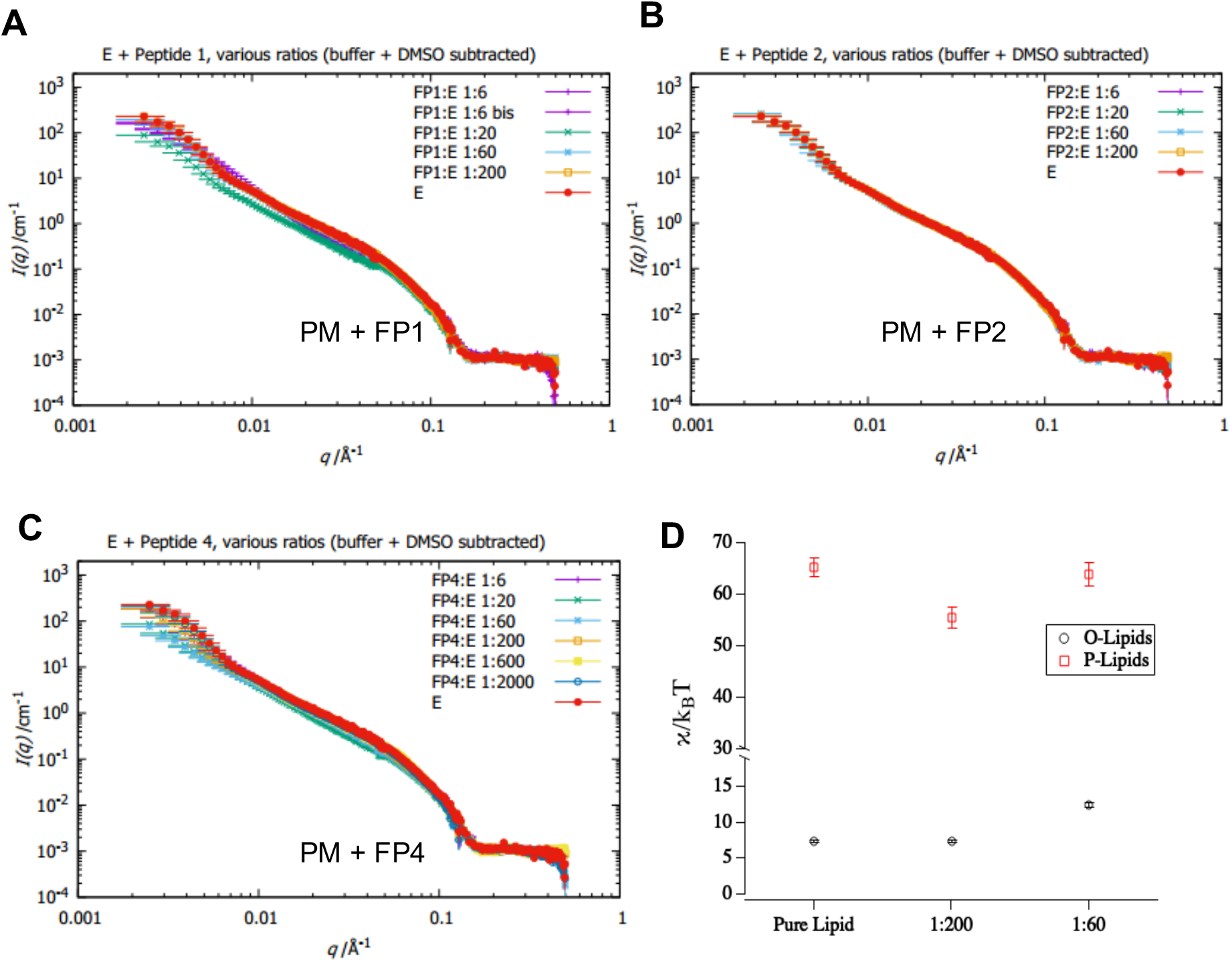
**A, B, C** SANS analysis of samples consisting of **A** FP1, **B** FP2, and **C** FP4 and extruded vesicles of lipids without (“O”) and with (“E”) cholesterol and sphingomyelin, in Kratky plot (I(q)q² vs q). The caption indicates the mole ratio lipid:peptide. Experimentally, mixtures were prepared starting from the highest concentration of peptides, and more diluted compositions were only explored as long as effects were seen in the SANS data, which is why for FP3 only the highest ratio is available (no effect in the investigated SANS q-range, least effective peptide), while for FP4 effects were visible down to low concentrations of peptide (most effective peptide). Overall effects are relatively small by SANS (preservation of membrane content) compared to macroscopic observation (large aggregates formation). FP3 and to a lesser extent FP3 have almost no effect. FP1 and mostly FP4 show the largest effects. For FP1, the change is in line with disk formation (constant amount of membrane but decreasing number of large vesicles), suggesting membrane perforation and at the largest peptide concentration stabilization of disks. For FP4, the increase of size observed at low q suggests binding and fusion between vesicles. **D** NSE-derived bending rigidities of PM liposomes in the presence of FP1. Addition of small amounts of FP1 leads to a slight softening of the membrane. Addition of larger amounts of FP1 leads to an apparent increase of rigidity for “O” lipids lacking cholesterol, which is probably due to the formation of multilamellar structures. Data from IN15 at ILL.

The addition of any FP led to turbidity of the SUV solutions. The extent of flocculation increased from FP3 to FP2 to FP1 to FP4, with FP3 being barely turbid and FP4 the most turbid. **Figure 3 A, B, C** show the SANS patterns of different FP bound to PM liposomes. Similar effects were also observed in the absence of cholesterol, as well as with ERGIC membranes (shown in **Figure S6**), indicating that the cholesterol did not affect FP action on the liposomes. Although due to the SUV heterogeneity, it was not possible to describe their geometry analytically, differences in scattering provided molecular insights about the FP action on the SUVs (**Figure S6 A**). Given the changes observed at low and at mid q, FP1 appears to disrupt the SUVs, followed by liposome association. The presence of FP2 and FP3 does not lead to scattering changes in the liposome SANS profiles, although some aggregation is observed. The effect of FP4 was different: the onset of the plateau was washed out, thus demonstrating a significant growth of the SUV structure size. This could be interpreted as an FP4-induced clustering of liposomes.

These results agree with the SNR data and indicate that FP1 buries into the lipid membrane. FP4, on the other hand, interacts primarily with the lipid head groups and is able to bind both the plasma and the viral (ERGIC-like) membranes simultaneously, thus form a bridge between different liposomes. In previous studies, fluorescence experiments showed that a section of FP4 has a fusogenic effect on SUVs that is enhanced in the presence of cholesterol (15).

### FP1 increases membrane rigidity in the absence of cholesterol

It is commonly known that membrane dynamics exhibit a multitude of different motions. The influence of FP1 on lipid membranes was investigated by NSE measurements on the nanosecond time and nanometer length scales.

Contrary to the expectation of a more fluid membrane with lower membrane rigidity, the addition of FP1 at 1:60 peptide:lipid molar ratio to “O”-SUVs (lacking cholesterol) in the absence of calcium leads mesoscopically to a significantly stiffer membrane (**Figures 3 D** and **S7**). Although possibly due to liposome association induced by FP1, this observation could be equivalent to the increase in membrane ordering previously detected from the electron spin resonance (ESR) signal of spin labels attached to lipids (19). Moreover, addition of peptide to (cholesterol-rich) PM-SUVs had hardly any effect, which is likely due to their bilayers’ inherent stiffness. The increase in bending rigidity at higher FP1 concentrations can be explained by the peptide inserting into the lipid acyl chains, and not by interactions with cholesterol.

### FP1 and FP2 binding affect the fluidity of the plasma membrane

The impact of FP1 and FP2 on the lipid membrane local dynamics on the picosecond time and Angstrom length scales was also probed by QENS (**Figure 4** and **S8**). At the sub-nanosecond time scale, lipid chain motions, including defect motions and rotational diffusion about the lipid molecular axis (occurring on the 10^−11^ s time scale) predominate. The 10^−10^ – 10^−9^ s time frame is also characterized by the presence of vibrational motions of the lipid molecules across the membrane (31, 32). Lateral lipid diffusion, rotational and flip flop motions of the lipid head occur at longer times (few ns), while collective oscillation of the whole membrane, providing information on the membrane roughness, is detectable on the 0.1 – 1 s time scale.

**Figure 4.**
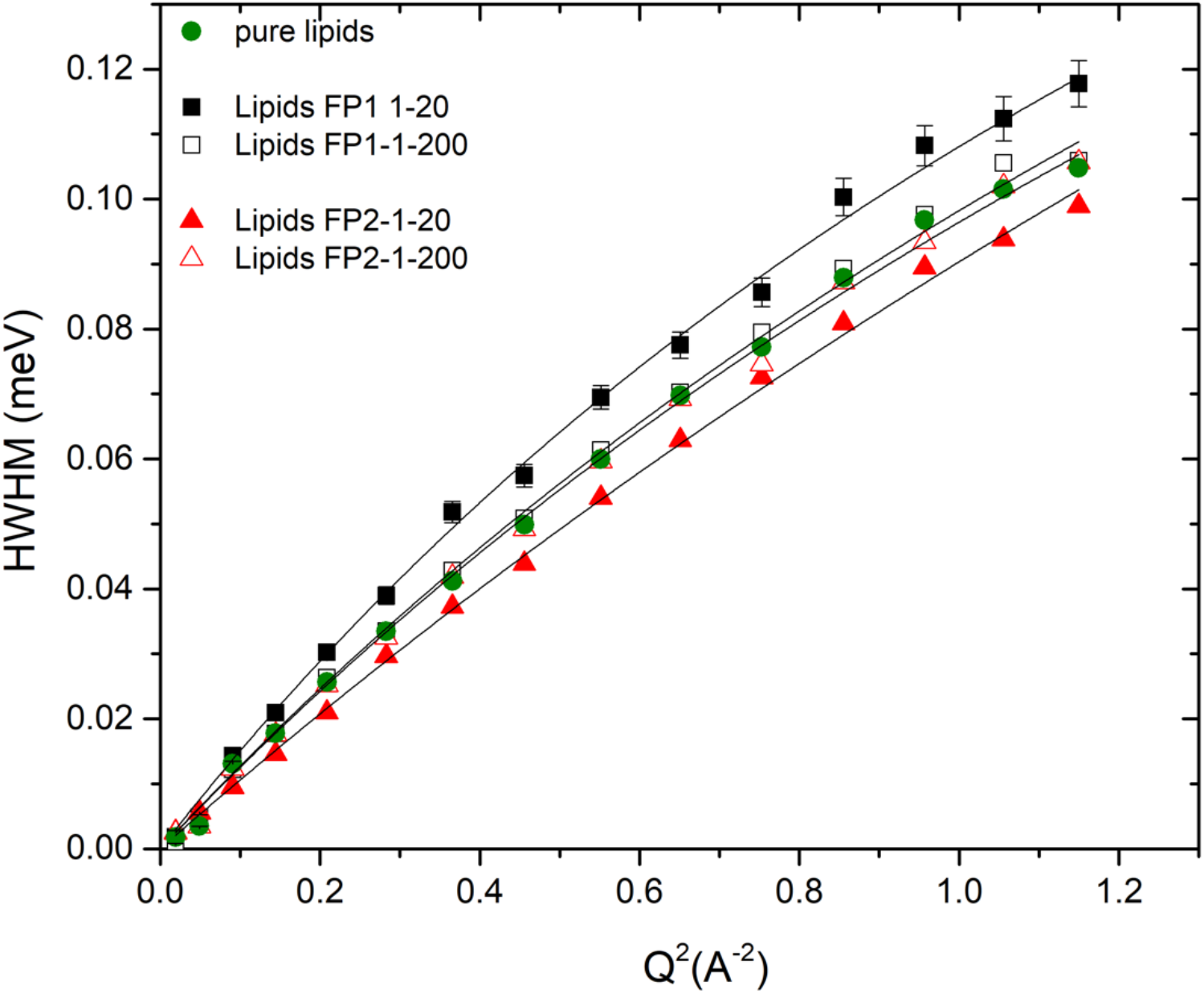
QENS- data derived half width at the half maximum (HWHM), of the Lorentzian function plotted versus q^2^ for the pure lipids, and lipids-peptide complex at high (1:20) and low concentration (1:200) FP1 and FP2 peptides. The straight line represents the fit to the data performed using the jump diffusion model as described in SI. The observed scattering signal arises mainly from the lipid membrane and represents an ensemble-average over the diffusive motions of the prevailing hydrogens atoms. The observed differences in the measured averaged signal are significant as per the diagonal elements of the covariance matrix of the fits (i.e., neglecting correlated errors). Data from IN5 at ILL.

In the jump diffusion model (33, 34), the QENS data describes the local dynamics of the hydrogen atoms throughout a residential time spent at a specific site and the diffusion coefficients to move between sites. The inferred parameters are reported in **Table 1**. The lack of a significant effect upon addition of FP1 and FP2 at both 1:20 and 1:200 peptide:lipid molar ratios, is observed for the diffusive component describing the fast dynamics at short range (10^−12^ s (or ps) time scale and few Å length scale) (see **Figure S8**). A strikingly different behavior is detected for the slow diffusion dynamics of the whole lipid acyl chain (time scale ≈ 100 ps). While no relevant effect on the membrane dynamics at the low peptide concentration (1:200) is observed, at higher peptide concentration (1:20), the dynamics of the FP-lipid samples show a significant deviation in comparison to pure lipids (**Figure 4**). A 25% faster diffusion coefficient of the lipid tails is measured in the presence of FP1, while in the case of FP2 the diffusion coefficient is reduced by a quarter.

**Table 1.** Diffusion coefficients and residence times inferred from the jump diffusion model

|  | **D (\*10<sup>-5</sup> cm<sup>2</sup>/s)** | **t (ps)** |
| --- | --- | --- |
| *Pure lipids* | 1.93+/-0.03 | 1.7+/- 0.1 |
| *Lipids+FP1 (1:200)* | 1.97+/-0.03 | 1.65 +/- 0.1 |
| *Lipids+FP1 (1:20)* | 2.36+/- 0.04 | 1.9 +/- 0.1 |
| *Lipids+FP2 (1:200)* | 1.97+/- 0.02 | 1.65 +/- 0.1 |
| *Lipids+FP2 (1:20)* | 1.62+/- 0.02 | 1.3 +/- 0.1 |

Results suggest that for FP1 peptides in the absence of calcium, the altered dynamics is certainly related to a change towards increased disorder in the local environment of the lipid tails, possibly due to the manner FP1 binds into the acyl region. The fastening of the tail fluctuations is in agreement with molecular dynamics simulations from Larsson and Kasson (35). Membrane ordering previously observed by ESR was in the presence of calcium (18, 19). The hypothesis that FP1 induced fusion between viral and host membrane could also provoke a lipid ejection and therefore a change in the local lipids tail order and spatial constraint, can justify not only the increase of the diffusion coefficient (more free space to explore among sites) but also the longer residential time since, in this scenario, atom-atom interactions are not affected by geometrical constraints.

FP2 peptides, which mostly lie down on the membrane surface, induce a moderate diffusion across the membrane (confined vertical fluctuation), which is coupled to the lipid tails rigidity. These changes in lipid tails movements could possibly be indirectly due to the dehydration effect of FP2 on the lipid head groups, as observed by SNR.

The increase in fluidity in the presence of FP1 and decrease in the presence of FP2 observed by QENS is consistent with the decreased tail density seen in the vertical density profiles in the presence of FP1 and increased density in the presence of FP2. A higher density would simply reduce the flexibility of the tails.

## DISCUSSION

SARS-CoV-2 fusion peptides FP1, FP2 and FP4 behave in a striking different manner in the presence of membranes. Fusogenic activity does not necessarily have to occur through membrane disruption but can be simply achieved by the close association of two membranes. By binding with greater affinity to the lipid head groups, FP4 is better able than FP2 to bridge two bilayers together. In either case, the observed dehydration of the lipid head groups would promote membrane fusion. Dehydration was for example found to be the rate-limiting step in SNARE-assisted membrane fusion (36). The kinetic barrier to hemifusion is generally estimated to exceed 50 kcal/mol (9). The binding free energy for association of a single FP4 with a lipid bilayer is ∼10 kcal/mol, so the pull on a Spike trimer consisting of three FP4 would be ∼ 30 kcal/mol. The kinetic barrier can therefore be overcome upon binding two Spike trimers.

FP1 disrupts the membrane as the peptide penetrates into the hydrophobic acyl region, and increases the dynamics of the lipid tails on a picosecond time and Angstrom length scale, which we here probed by QENS. This increase in alkyl chain fluidity is consistent with previous observations of lipid disordering that might be associated with a subsequent weakening of lipid membranes (e.g. more prone to rupture), as an effect of lipid-FP interactions. This weakening of lipid membranes has previously been interpreted as a first step towards viral penetration into the host cell (37). On the nanosecond time scale probed by NSE at low FP1 concentrations in the absence of calcium, an increase in membrane flexibility is more pronounced in the presence of cholesterol-rich PM. A phenomenon also observed by QENS. However, at increased FP1 concentrations, NSE indicates the SUVs membrane rigidity increases, which could be analogous to the increase in local membrane ordering in multi-lamellar vesicles identified by electron spin resonance (ESR) spectroscopy (18, 19). These effects could possibly be due to the FP1-driven membrane stacking in the absence of calcium, which was observed in the SANS data. A membrane consisting of several bilayers is simply more rigid than a single bilayer.

Free Ca^2+^ concentration varies widely depending on its cellular location. At the plasma membrane, outside the cell, [free Ca^2+^] ∼ 2mM, while in the early and late endosomes [free Ca^2+^] drops to ∼ 0.3μM (11). The affinity of SARS-CoV-2 FP1-2 and the related SARS-CoV-1 FP1 to calcium (∼30 μM), determined by ITC (19), would suggest that calcium would not be bound to the fusion peptide in the endosome. As with the calcium-dependence in lipid membrane ordering observed by ESR, our SNR data show a clear effect of the cation on SARS-CoV-2 FP1 and FP2 PM binding. The binding efficiency of the latter increases, but more intriguingly, calcium drastically alters the orientation of bound FP1 in the PM. The intracellular calcium levels may therefore provide an indication to where the viral and host membranes fuse during SARS-CoV-2 infection.

Depending on cell type, the SARS-CoV-2 virus may enter the host cell through the plasma membrane (10), but can also travel through the endosomal pathway (38). Although the virus may have associated with the host at the PM, it is unclear where actual S2’-mediated membrane fusion occur. It may even occur in the late endosome (39). Our structural data show that at the PM, in the presence of calcium, the FP1 peptide would penetrate the PM and form a transmembrane peptide across the bilayer. Subsequently as free calcium levels drop in the endosome (11), FP1 re-positions itself to the host lipid leaflet in contact with the viral membrane. Its shallower position then enables it, like FP4, to function as a bridge between the host and the viral membrane, as was shown by SANS.

Two competing models have been put forward to explain protein-driven membrane fusion (reviewed in Lindau and Almers (40)). In the “proximity” model, two membranes are brought together by a fusion peptide, like FP4, and the juxtaposed leaflets of each bilayer merge and their lipids mix. In the “fusion pore” model, initially proposed by Pfenninger (41), membrane hemifusion is initiated by the creation of an aqueous channel, which is tentatively observed for FP1 in the presence of calcium at the PM. Our structural data from SANS show FP1 can associate and consequently possibly puncture through the viral membrane, thereby fusion initiation points occur in both the PM and viral membranes. The initiation points would subsequently expand to allow lipids to travel along the amphipathic fusion peptide and diffuse between the two membranes (**Figure 5 A, B**). Importantly, when the influenza haemagglutinin transmembrane domain was replaced by glycolipid anchors, the transfer of fusing membranes occurred at the same rate, even though the viral pore formed solely in the eukaryotic cell membrane (42), so fusion initiation point formation in the viral membrane may not be necessary. By bridging the membranes together, the subsequent role of FP4 and FP1 (after Ca^2+^ removal), would be to bind and dehydrate the two membranes in order to drive the expansion of a hemifusion diaphragm formed between the viral RNA and cellular cytoplasm (**Figure 5 C, D**). The rest of the S2’ domain, due to its extra-membrane bulk, would be excluded from the interface and be expelled from the contact area formed between the viral and host membranes.

**Figure 5.**
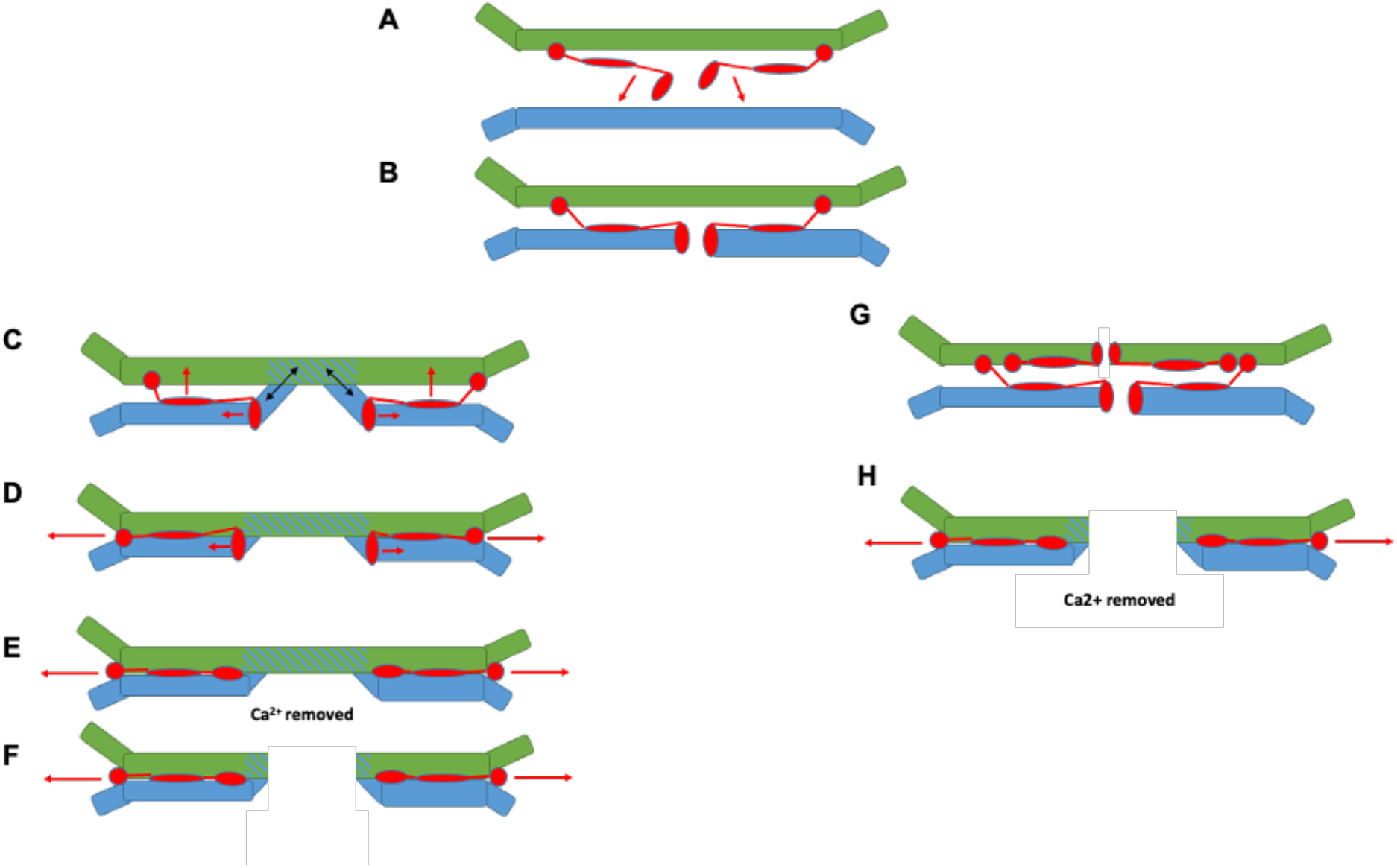
Proposed fusion mechanism between SARS-CoV-2 and eukaryotic host membrane. The viral membrane bilayer is colored green, the eukaryotic host membrane in blue, and the S2’ protein is in red. The direction of the protein is drawn with red arrows, while the direction of the lipids is drawn with black arrows. FP1and FP4 are represented as ovals, and the structured S2’ protein as a circle (attached to the viral membrane). **A** FP1 forms a fusion initiation point on binding the host membrane. **B** The initiation point enlarges provoking lipids mixing between the viral and host membrane, leading to the growth of a hemifusion diaphragm. **C** FP4 bridges the two membranes together thereby facilitating the fusion of the two membranes into a single bilayer. Moreover, the two membranes coming together exclude the folded S2’ from the growing synapse. **D, E, F** A hemifusion diaphragm is formed and in the endosome, lower free calcium concentrations lead to FP1 orienting itself like FP4, thereby providing further contact between the two membranes. **G** It is also possible that FP1 initiation points may form on both the viral and the Plasma membrane. **H** The two membranes form a pore, and as in **F**, the pore expands as the Spike protein is excluded by the two membranes coming together due to the bridge function encoded in the Spike fusion peptides.

In conclusion, our multi-method approach based on time-averaged and energy-resolved neutron scattering methods applied to a well parametrized model of the protein-host membrane interaction, reveals that different segments of the SARS-CoV-2 Spike protein assume different functions in the initiation of viral infection. Our data are of interest not only in the context of the current Covid-19 pandemic, but also provide a powerful interdisciplinary framework for future investigations of eukaryotic and viral fusion mechanisms.

## MATERIALS AND METHODS

### Fusion Peptides

The fusion peptides were synthesized and purified by GenScript (Amsterdam, The Netherlands). Stock solutions of each peptide in DMSO (dimethyl sulfoxide) were used for all the experiments reported here. The following peptides were investigated (**Table S1**):

> FP1 (SARS-CoV-2 816-SFIEDLLFNKVTLADAGFIKQY-837),
>
> FP2 (SARS-CoV-2 835-KQYGDCLGDIAARDLICAQKFN-856),
>
> FP3 (SARS-CoV-2 854-KFNGLTVLPPLLTDEMIAQYT-874) and
>
> FP4 (SARS-CoV-2 885-GWTFGAGAALQIPFAMQMAYRFNGI-909).

### *Pichia pastoris* cell culture

*P. pastoris* cells were cultured in the Deuteration facility of the Institut Laue-Langevin (D-Lab, ILL), Grenoble, France, using the protocol described elsewhere (26). Cells were grown in flasks at 30 °C using a basal salt medium (BSM) as the minimal medium at pH 6.0 (*P. pastoris* fermentation process guidelines, Invitrogen, United States) containing either 20 g.L^−1^ glycerol in H_2_O or glycerol-d8 (Euriso-Top, France) in D_2_O. Cells upon entering the exponential phase at an OD of 600 were harvested by centrifugation and frozen at -80 C. Lipid extraction and separation was performed in the L-Lab facility within the Partnership for soft condensed Matter (PSCM) at the Institut Laue-Langevin, Grenoble, France (ILL). The exact methodology is described in the **SI Methods**.

### Natural lipid monolayers formulation

Biomimetic membranes in the form of monolayers were prepared from phosphatidylcholine (PC), phosphatidylethanolamine (PE) and phosphatidylserine (PS) extracted and purified from perdeuterated and hydrogenous *P. pastoris* cell cultures (SI methods). Hydrogenous cholesterol and egg yolk sphingomyelin were purchased in powder form from Sigma Aldrich. Deuterated cholesterol was obtained from the National Deuteration Facility (NDF) in ANSTO (Australia). For biomimetic PM, the composition in molar ratio was PC 0.2, PE 0.11, PS 0.06, cholesterol 0.5, sphingomyelin 0.13 (43) (**Table S2**).

### Synthetic lipid vesicles for solution and for supported lipid bilayer (SLB) studies

1-palmitoyl-2-oleoyl-sn-glycero-3-phosphocholine (POPC), 1-palmitoyl-2-oleoyl-sn-glycero -3-phosphoserine (POPS) and cholesterol were purchased from Avanti Polar Lipids and used without further purification. Small unilamellar vesicles (SUV) were prepared by dissolving POPC, POPS and cholesterol in chloroform, mixing according to desired membrane composition, (i.e. 3:1:1 molar ratio, respectively, as previously described in the Fusion Peptide literature (17, 18)), dried under gentle Argon flow and placed in vacuum overnight to ensure evaporation of all solvent. The resulting lipid films were rehydrated at room temperature in buffer at up to 1 mg·mL^−1^ lipid concentration, and vortexed to fully suspend vesicles. Immediately before use for supported lipid bilayer (SLB) formation, the suspension was tip sonicated for 5 min at pulses of 1 s on/off to produce a visually clear solution of SUVs. SLB formed from vesicle fusion on the surface.

### Small unilamellar vesicles (SUVs) for neutron spectroscopy measurements

Lipids were dissolved in chloroform, dried under gentle Argon flow and placed in vacuum overnight to ensure evaporation of all solvent. The resulting lipid films were dissolved in D_2_O at up to 50 mg/ml lipid concentration. After several cycles of sonication, liposomes were prepared by filtration through a 500 nm pore-size Millipore filter in order to remove aggregates. FP1 and FP2 peptides, which were dissolved in 50 % (v/v %) of DMSO D_2_O buffer solution, were then mixed with the lipids, and after further tip sonication, SUVs were prepared by extrusion through a 100 nm pore-size membrane, using an Avanti® Mini-Extruder. The outcome of the extrusion was checked by Dynamic Light Scattering (Malvern Nanosizer at the PSCM).

### Circular Dichroism (CD)

Measurements were performed on a Jasco J-810 spectropolarimeter. Spectra with a band width of 2 nm, an acquisition speed of 50 nm.min^-1^ and an integration time of 1 s were collected using 0.1 cm pathlength cuvettes. Peptides at a final concentration of 29 μM in 10 mM phosphate buffer pH = 7, in the presence and absence of 100 nm pore size extruded SUVs (1:100 peptide:lipid ratio) were analyzed. Blanks, i.e. the phosphate buffer spectrum, as well as the purely liposome spectrum, were also measured. Raw millidegree data were converted to molar ellipticity.

### Lipid monolayer and Langmuir trough experiments

A Langmuir trough, available at the PSCM, with a total area of 166.4 cm^2^ equipped with two dependent barriers (Kibron, Helsinki, Finland) was used to measure the surface pressure (Π) of lipid monolayers and the increase in pressure (ΔII) observed after the injection of the peptides in the bulk phase. The variation of surface pressure was recorded using a Wilhelmy plate made of filter paper. Temperature was maintained at 21.5 ± 0.5 °C. Further experimental details are given in the **SI Methods**.

### Brewster Angle Microscopy (BAM)

*In situ* visualization of the morphology of Langmuir monolayers at the air/water interface was performed using a BAM Nanofilm EP3 (Accurion, GmgH, Goettigen, Germany) available at the PSCM. The instrument was equipped with a 50 mW laser emitting p-polarized light at a wavelength 532 nm directly onto the air/water interface at the Brewster angle (53.1°) through a 10x magnification objective and a polarizer (44). The reflected light is captured by a CCD camera. The spatial resolution was 2μm and the field of view 200 × 200 μm^2^.

### Specular neutron reflectometry

Experiments were performed on the time-of-flight reflectometer FIGARO at the ILL. Two different angles of incidence (θ_1_ = 0.6° and θ_2_ = 3.7°) and a wavelength resolution of 7% dλ/λ, yielding a momentum transfer, q=(4π/λ) sinθ, range from 0.007 to 0.25 A^-1^ (the upper limit being limited by sample background) were used to perform the measurements and investigate the structure of the lipid monolayers and bilayers upon peptide interaction. Lipid monolayers were prepared in a Langmuir trough at a surface pressure of Π = 23 ± 1 mN m^-1^. After SNR measurements of the lipid monolayer, peptides were injected under the monolayer by a Hamilton syringe to a final bulk concentration of 3µM. Two different isotopic solvent contrasts (100% D_2_O and 8.1% D_2_O (v/v %) respectively) were used for the characterization. For SNR experiments on SLB, solid/liquid flow cells available at the ILL with polished silicon crystals (111) with a surface area of 5×8 cm^2^ were used. Substrate surfaces were characterized in 2 different isotopic solvent contrasts, (0% and 100% D_2_O), before SLB formation. The membranes were subsequently characterized in at least 3 isotopic solvent contrasts (100% H_2_O, 100% D_2_O and 38% D_2_O buffers). The SNR data were reduced and normalized using COSMOS (45). Subsequent data analysis was performed using AuroreNR (46) and Motofit software (47) (**SI Methods**).

### Small-angle neutron scattering experiments

SANS measurements were carried out on D22 at the ILL, using Hellma quartz 120-QS cells of 1 mm pathway. Samples were measured over a q-range of 2.5 10^−3^ to 0.6 Å^-1^ at a single wavelength of 6.0 Å (FWHM 10 %) with 3 sample-to-detector distances of 1.5 m, 5.6 and 17.6 m. Absolute scale was obtained from the flux using the attenuated direct beam. Data correction was performed using Grasp, accounting for transmission, flat field, detector noise (measurement of boron carbide absorber); the contribution from the solvent was subtracted.

### Quasi elastic neutron scattering

Experiments were performed using the high-resolution direct geometry Time-of-Flight (ToF) spectrometer IN5 at the ILL. In order to investigate molecular motions of the lipids, two different experimental configurations were studied: at neutron wavelength 10 Å and energy resolution of 10 µeV FWHM (corresponding to a time resolution of ! 70 ps and a q-range between 0.2 and 1.2 Å^−1^), and at 5 Å and 70 µeV FWHM (corresponding to a time resolution of ! 10 ps and a q-range between 0.27 and 2.14 Å^−1^). All measurements were performed at 300 K. The spectra were corrected for the detector efficiency, subtracted by the background and normalized following the standard procedures by a vanadium spectrum, by using the sofware package lamp (https://code.ill.fr/scientific-software/lamp). The QENS data were analyzed using the DAVE program (https://ncnr.nist.gov/dave), with a delta function mimicking the elastic part and a Lorentzian line-width accounting for the ensemble-averaged contribution from the peptides and lipids.

### Neutron spin-echo experiments

Measurements were performed on the instrument IN15 at ILL using neutron wavelengths from 8 to 13.5 Å covering a q-range from 0.03 to 0.14 1/Å and Fourier times up to 477 ns. Data were analyzed using the Zilman-Granek model (48) (see **SI Methods**). The samples measured were at a lipid concentration of 10 mg/ml.

## Acknowledgements

The authors thank the ILL for awarding beam-time: **SNR** and **SANS**, DOI: 10.5291/ILL-DATA.DIR-215, and **QENS** and **NSE**, doi:10.5291/ILL-DATA.DIR-211) and use of support facilities at the D-Lab and Partnership for Soft Condensed Matter (PSCM). Financial support for consumables was also provided by the Science and Technology Facilities Council (U.K.). N.R.Z was supported by Wellcome Trust grant WT 207455/Z/17/Z. Part of the lipid extraction activity was funded by the ANR/NSF-PIRE project REACT (Research and Education in Active Coatings Technologies for Human Health). The National Deuteration Facility in Australia is partly funded by The National Collaborative Research Infrastructure Strategy (NCRIS), an Australian Government initiative.

We gratefully acknowledge M. Jourdan and J. Dejeu (Université Grenoble Alpes) for access to the CD instrument. J. Carrascosa for help in the analysis of the SNR data collected, and, Prof. E. Guzman, Prof. P. Luzio, Prof. D. Owen and Dr. J. Zaccai for a critical reading of this manuscript.

## Author contributions

A.M and N.R.Z. designed research; A.S, A.M, D.R, F.N, G.C., O.M., S.P., and T.S. performed research; A.S, A.M, D.R, F.N, G.F., N.R.Z., O.M., S.P., and T.S. analyzed data; K.C.B., M.H., R.A.S., T.D. and V. L. contributed new reagents; all authors discussed and reviewed the results; and N.R.Z and A.M. wrote the manuscript with input from all authors.

