## Supplementary Information for "Strikingly different roles of SARS-CoV-2 fusion peptides uncovered by neutron scattering"

**This PDF file includes:**

Supplementary methods
Tables S1 to S6
Figures S1 to S7
SI References

**SUPPLEMENTARY METHODS**

**Natural lipid extraction and purification for lipid monolayer studies**

Biomimetic membranes were prepared from phospholipids mixtures extracted and purified from perdeuterated and hydrogenous *P. pastoris* biomasses. Harvested cells were suspended into 10 ml deionized water and lysed by probe sonication on an ice bath for 3×5 min with 30 s intervals, 20% duty cycle. The resulting cell lysate was poured into boiling ethanol containing 1% butylated hydroxytoluene (BHT) followed by vigorous stirring in order to denature lipases. The total lipid mixtures were then extracted according to the method of Folch et al. (1), followed by evaporation of the organic phase under a $N_2$ stream and their final reconstitution in $CHCl_3$.

Purification of the various classes of phospholipid mixtures containing molecular species of mixed acyl chain lengths was achieved through sequential purifications, first by passing the lipid extracts through an amino-bonded solid-phase extraction column followed by purification through a diol-modified silica stationary phase column coupled to a High-performance Liquid Chromatography-Evaporative light scattering detector (HPLC-ELSD) Agilent 1260, United Kingdom) system. The mobile phase employed was a gradient between solvent A ($CHCl_3$/$CH_3OH$/$NH_4OH$, 80:20.5:0.5, v/v) and solvent B ($CHCl_3$/$CH_3OH$/$H_2O$/$NH_4OH$, 60:35:5.5:0.5, v/v)(2). TLC analysis was carried out on a High-Performance Thin-Layer Chromatography (HPTLC) system (CAMAG, Muttenz, Switzerland) to assess the identity and purity of each of the purified classes. Fatty acid compositions of such purified mixtures were measured by Gas Chromatography-Flame Ionization Detection (GC-FID).

**Lipid monolayer fabrication and determination of the binding affinity**

A Langmuir trough with a maximum total area of 166.4 $cm^2$ equipped with two dependent barriers (Kibron, Helsinki, Finland) was used to measure the surface pressure ($\Pi$) – area isotherms of lipid monolayers (See **Fig. S1 A**). $\Pi$ is recorded using a Wilhelmy plate made of filter paper (Whatman CHR1 chromatography paper).

A home-made trough with a total area of 106 $cm^2$ were used to determine the peptides binding affinity. After careful cleaning, the trough was filled with the buffer (HEPES 5mM, NaCl 150mM), and a 0.2 $mg \cdot mL^{-1}$ lipid solution in chloroform was spread over the subphase using a Hamilton micro-syringe to achieve an initial surface pressure of 22 mN $m^{-1}$, to mimic the outer leaflet of the plasma membrane. The subphase temperature was maintained at 21.5 ± 0.5 °C by making thermostated water flow through jackets at the bottom of the trough. After the evaporation of the chloroform and the stabilization of the monolayer, peptides were injected underneath the lipid monolayer in the bulk phase. The binding constant of the peptides with the lipid monolayer was calculated by measuring the increase in pressure, $\Delta\Pi = \Pi_\infty - \Pi_0$, observed after the injection of the peptides in the bulk phase, $\Pi_0$ = 22±1 mN.$m^{-1}$, until the surface pressure reached a plateau, $\Pi_\infty$. The differences in the surface pressure $\Delta\Pi$, which depend on the amount of peptide injected, were normalized with respect to the maximum value of difference obtained ($\Delta\Pi_{max}$). The obtained $\Delta\Pi/\Delta\Pi_{max}$ versus peptide concentration profiles were analyzed with a Langmuir adsorption model based on an adsorption equilibrium equation and mass balance between the monolayer and the FPs, assuming that the degree of normalized surface pressure increase is proportional only to the coverage fraction of peptides interacting with the monolayer φ:

$$\varphi = \frac{\Delta\Pi}{\Delta\Pi_{max}} = \frac{C^n}{C^n + k_d} \qquad \textbf{(S1)}$$

where C is the bulk concentration of the peptide, n is the Hill coefficient and $k_d$ is a dissociation constant (3).

**Specular Neutron reflectivity (SNR)**

***Definition.*** SNR elucidates the structure and composition of monolayers and bilayers in the direction perpendicular to the plane of the interface. A 1D profile of the reflectivity (R), defined as the ratio of neutrons scattered from the interface over the incident intensity of the neutron beam, is measured in specular conditions (i.e., the incident angle of the neutron beam is equal to the reflected angle, denoted as θ) as a function of the momentum transfer vector q normal to the interface (defined as $q = 4\pi \sin\theta / \lambda$, where λ is the wavelength of the neutron beam). The measured reflectivity can be linked to a plane-averaged scattering length density (SLD) profile perpendicular to the interface. SLD measures the coherent scattering cross-section of the molecular species that constitutes each interfacial layer, and is linked to their chemical composition and molecular volume, V by:

$$SLD = \sum_i b_i . n_i \qquad \textbf{(S2)}$$

where $n_i$ is a number density and $b_i$ is the scattering length density of each molecular species.

***Data acquisition.*** SNR measurements were performed on FIGARO, a time-of-flight reflectometer, at the Institut Laue-Langevin (Grenoble, France). Two angles of incidence θ (0.62° and 3.8°, for Langmuir monolayers and 0.8° and 3.2°, for solid supported lipid bilayers, respectively) and a wavelength resolution of 7% dλ/λ were used yielding a momentum transfer range of $0.005\ 01 \leq q \leq 0.25$ Å$^{-1}$ and a residual background reflectivity of $R \sim 10^{-7}$. The raw time-of-flight experimental data at these two angles of incidence were calibrated with respect to the incident wavelength distribution and the efficiency of the detector yielding the resulting R(q) profile using COSMOS (4).

Measurements were performed in 5mM HEPES, 150mM NaCl buffer (pH 7.1) in different isotopic contrasts, such as $H_2O$, $D_2O$, 8.1% v/v $D_2O$ (denominated air-contrast matched water, ACMW, which has an SLD of zero), and 38% v/v $D_2O$ (in order to match the silicon SLD, i.e. SLD = 2.07 $10^{-6}$ Å$^{-1}$).

***Data modelling*** was performed by minimizing the difference between the experimental data points and the calculated reflectivity profile. The latter was obtained by a model consisting of multi-layers of constant SLD using the Parratt's recursive method, with an error function connecting adjacent layers. Data analysis was performed using constrains between layer parameters (thickness, roughness and degree of hydration) and simultaneous co-refinement of all data sets obtained with a global minimization of a least-squares function $\chi^2$ was done to reduce the ambiguity in the modelling by using both AuroreNR (5) and Motofit (6) packages. Following (7), the area-per-molecule of lipidic components was fixed to be the same in each layer, $APM = {}^{b_i}/_{SLD_i t_i f_i}$ (with i=aliphatic tails or head-groups). In presence of the peptides, in each layer (i.e., tails and heads for mimicking membrane leaflet) the SLD and the volume fraction $f$ follows: $\rho_{model} = f_{solvent}.\rho_{solvent} + f_{lipid}.\rho_{lipid} + f_{peptide}.\rho_{peptide}$; where $f_{solvent} + f_{lipid} + f_{peptide} = 1$. All the parameters used in the fitting process and results are tabulated in **Tables S3**, **S4** and **S5**.

In the case of solid-supported bilayers, solid/liquid flow cells with polished silicon crystals (111) with a surface area of 5×8 cm$^2$ were used. Substrate surfaces were characterized in 2 different isotopic solvent contrasts, (100% $H_2O$ and 100% $D_2O$), before SLB formation. Lipid bilayers were deposited on silicon crystals by vesicle fusion. The presence of bilayers was confirmed by simultaneously fitting SNR profiles measured in 3 isotopic contrasts (see **Figures S4, S5**). Where possible, the bilayer structure was constrained to be symmetric, so that the inner and outer leaflets would be equivalent. The head group molecular volumes and thicknesses were fixed to values obtained from X-ray diffraction measurements (8), while the acyl tail region, had a molar averaged SLD value (see **Table S4**). The resulting model consists of four layers describing the inner and outer head groups, the lipid tail region and a layer of water between the $SiO_2$ layer and the inner lipid leaflet. The acyl tail region had a low water content for all the bilayers, confirming a lipid coverage of at least 85%. A 3 to 6 Å thick water layer is also present between the bilayer and the silicon crystal. The average area per molecule obtained of 53 to 59 Å$^2$ and the total thickness of 47 Å agree with similar systems reported in the literature (9).

**Small Angle Neutron Scattering (SANS)**

Solution small-angle neutron scattering (SANS) probes the shape, the spatial distribution and the interactions between solute particles by detecting the radiation scattered by the sample as a function of momentum transfer q previously defined.

The differential scattering cross-section of a SANS experiment can be expressed as (10, 11),

$$\frac{d\sigma}{d\Omega}(q) = n(\Delta\rho)^2 V_{part}^2 P(q) S(q) \qquad \textbf{(S3)}$$

where n is the particle number density, $\Delta\rho$ is the difference in scattering contrast between the solvent and the particles, and $V_{part}$ is the volume of a single particle. The term P(q) is referred to as the particle form factor, determined by the protein conformation (11). The structure factor, S(q) (for details, see e.g. Refs. (12-14) ), characterizes the correlations and thus the interaction between proteins in solution, and the position of the principal peak can be related to a characteristic distance $\xi=2\pi/q$ in the sample. In the low-q region, the relation $I(q) / q^{-D}$ can be used, where the fractal dimension D of the system provides additional structural information on the system in question (14-16). Here, the fractal dimension D describes how the mass of the respective fractals increases with distance from a given origin.

**Dynamics of biomacromolecules investigated by quasi-elastic neutron experiments.**

The dynamics of bio-macromolecules contributes to the thermodynamic stability of their functional states, covering a range of time ranging from femtoseconds for the vibrations of bonds, to milliseconds or seconds, for more complex processes that require a structural rearrangement. The dynamic landscape which corresponds to thermal energy is driven by movements at the picosecond to nanosecond timescale, which corresponds to the characteristic times of forces which stabilize the structures of bio-macromolecules, such as hydrogen bonds, "hydrophobic", Van der Waals and electrostatic interactions.

The study of dynamic processes at the microscopic level can be performed using neutron scattering and in particular, quasi-elastic neutron scattering (QENS), which offers the advantage of covering length scales (0.5-10 Å) and energies (0-200 MeV) corresponding to thermal fluctuations. The technique is sensitive to movements of hydrogen atoms, which are uniformly distributed within biological structures and represent nearly half of the atoms in them. On scales of time and space accessible with neutron scattering, the hydrogen atoms trace the movements of the chemical groups which they are linked to. QENS provides unique information useful for the characterization of the dynamical heterogeneity of biological macromolecules, from globular protein up to membrane complexes, in the time range $10^{-12}$ to $10^{-6}$s.

A quasi-elastic neutron experiment measures the incoherent dynamic structure factor $S_{inc}(q,\omega)$ which harbors all the dynamical information of the investigated system.

The dynamic structure factor can be written as follow

$$S(q,\omega) = A_o(q) \times \delta(\omega) + (1 - A_0(q) \times L(q,\omega) \quad \textbf{(S4)}$$

The first term represents the elastic part, accounting for particles whose motions cannot be resolved in the investigated time windows, while the second term define the quasi-elastic part, which identify the diffusion motions and can be described by a Lorentzian linewidth function.

Each linewidth depicts the average of active motions in the investigated time window. The q dependence of the half width at half maximum (HWHM) of the Lorentzian curve gives an insight into the nature of the diffusive motion. In particular, when representing the HWHM as function of $q^2$, the slope of the linear fit in the low q range provides information on the diffusion coefficient, since the motion reduces to continuous diffusion there, for which $\Gamma = Dq^2$. The QENS data was analyzed according to such an approach to get insights into motions of hydrogen atoms belonging to the lipids or the peptides. A combination of one Lorentzian and a delta function was required to optimize the fits. The buffer contribution was subtracted during the fitting procedure, feeding the program directly with the sample raw data and avoiding in such a way the necessity to fit a second Lorentzian curve.

We have found that the Lorentzian for translation is best fit to a random jump diffusion model, which considers the residence time τ0 for one site in a given network before jumping to another site (17, 18),

$$\Gamma^{J}_{trans}(Q) = \frac{D_{trans}Q^2}{1 + D_{trans}Q^2\tau_0} \quad \textbf{(S5)}$$

where the mean jump diffusion length L is defined as L= $\sqrt{6D_{trans}\tau_0}$ , and $D_{trans}$ is the translational diffusion coefficient between two sites. The jump diffusion model reduces to continuous diffusion in the low q range and tends to a plateau (∞) for higher q, with ∞= 1/0.

Measurements performed at 5 Å resolution, probing short dynamics (of the order of 5-10 ps), were not particularly conclusive. The fit to these spectra collected on IN5 results in an indistinguishable linewidth of the diffusive component, within the errors, for the different samples (**Figure S8**, which simply suggests that the probed fast dynamics at short range represents translational diffusion and is intrinsic of soft matter.

### NSE Theory

Data were analyzed using the Zilman-Granek model (19) but using a prefactor of 0.0069 instead of 0.025 (20-22), which arises from a combined bending-compression mode observed at the time and length scale of NSE (23) when calculating the bending rigidity k from the experimentally observed relaxation rate $\boldsymbol{\gamma}_{ZG}$. The normalized intermediate scattering function S(q,t)/S(q,0) has a stretched exponential shape,

$$S(q,t) / S(q,0) = \exp\{(-\gamma_{ZG} q^{3t})^{\frac{2}{3}}\} \quad \textbf{(S6)}$$

and $\boldsymbol{\gamma}_{ZG}$ is related to the bending rigidity by

$$\gamma_{ZG} = 0.0069 \Big/ \sqrt{\left(\frac{k_b T}{k}\right)\frac{k_b T}{\eta}} \qquad \textbf{(S7)}$$

where $k_B$ is the Boltzmann constant, T is temperature and $\eta$ is the viscosity of the solvent. While including a term for translational diffusion does affect the precise values of $\gamma_{ZG}$ obtained from the fits, the qualitative trend remains the same, whether it is included or not, which is why we chose to keep the analysis as simple as possible and present the fit results obtained without an explicit term for translational diffusion and a single free parameter $\gamma_{ZG}$ for each sample.

### SUPPLEMENTARY TABLES

| Peptide | Sequence | Number of AA | MW (Da) | pI | Net charge | Aliphatic index | Grand average of hydropathicity (GRAVY) |
|---|---|---|---|---|---|---|---|
| FP1 | **816-SFIEDLLFNKVTLADAGFIKQY-837** | 22 | 2538 | 4.56 | -1 | 110.91 | 0.368 |
| FP2 | **835-KQYGDCLGDIAARDLICAQKFN-856** | 22 | 2456 | 6.02 | 0 | 84.55 | -0.255 |
| FP3 | **854-KFNGLTVLPPLLTDEMIAQYT-874** | 21 | 2351 | 4.37 | -1 | 111.43 | 0.262 |
| FP4 | **885-GWTFGAGAALQIPFAMQMAYRFNGI-909** | 25 | 2723 | 8.75 | +1 | 66.80 | 0.516 |

**Table S1.** Physicochemical properties of the peptide investigated.

| Nomenclature | Lipid composition | Monolayer / bilayer | Experiments |
|---|---|---|---|
| **Synthetic model membrane** | POPC 0.6, POPS 0.2, cholesterol 0.2 (molar ratio) | Bilayer (SUV and SLB) | SLS, CD, NR |
| **PM** | PC 0.2, PE 0.11, PS 0.06, cholesterol 0.5, sphingomyelin 0.13 (molar ratio). | Monolayer | SNR, BAM, Langmuir titration |
| **"O" membrane (lacking cholesterol)** | PC 0.2, PE 0.11, PS 0.06, sphingomyelin 0.13 (molar ratio). | Bilayer (SUV) | SANS, QENS, NSE |
| **ERGIC membrane** | PC 0.2, PE 0.11, PS 0.06, cholesterol 0.2, sphingomyelin 0.13 (molar ratio) | Bilayer (SUV) | SANS |

*Nomenclature:

SLS: Static Light Scattering

CD: Circular Dicroism

SNR: Specular Neutron Reflectometry

BAM: Brewster Angle Microscopy

Langmuir titration -determination of binding affinity.

SANS: Small Angle Neutron Scattering

QENS: Quasi-elastic Neutron Scattering

NSE: Neutron Spin Eco.

**Table S2.** Lipid compositions used and experimental techniques

.

| Fixed Parameters | Plasma Membrane monolayer hydrogenous | Plasma Membrane monolayer deuterated |
|---|---|---|
| $V_{m\_heads}$ (Å$^3$) | 304.6 | 304.6 |
| $\Sigma b_{heads}$ ($10^{-5}$ Å) | 60.95 | 166.91 |
| $SLD_{heads}$ ($10^{-6}$ Å$^{-2}$) | 2.00 | 5.48 |
| $V_{m\_tails}$ (Å$^3$) | 764.1 | 803.2 |
| $\Sigma b_{tails}$ ($10^{-5}$ Å) | -1.40 | 425.62 |
| $SLD_{tails}$ ($10^{-6}$ Å$^{-2}$) | -0.02 | 5.30 |
| $f_{tails}$ (%) | 1 | |

**Table S3**. SLD, scattering length and molecular volume values used in **SNR** experiments performed on Langmuir lipid monolayers.

| Fixed Parameters | POPC:POPS:cholesterol bilayer |
|---|---|
| $V_{m\_heads}$ (Å$^3$) | 261 |
| $\Sigma b_{heads}$ ($10^{-5}$ Å) | 47.24 |
| $SLD_{heads}$ ($10^{-6}$ Å$^{-2}$) | 1.81 |
| $V_{m\_tails}$ (Å$^3$) | 864 |
| $\Sigma b_{tails}$ ($10^{-5}$ Å) | -16.43 |
| $SLD_{tails}$ ($10^{-6}$ Å$^{-2}$) | -0.19 |

**Table S4**. SLD, scattering length and molecular volume values used in the **SNR** experiments performed on solid-supported lipid bilayers.

| | PM monolayer | | | | PM monolayer + **FP1** | | | | | PM monolayer + **FP2** | | | | | PM monolayer + **FP4** | | | | |
|---|---|---|---|---|---|---|---|---|---|---|---|---|---|---|---|---|---|---|---|
| | *t* | $f_w$ | $f_{lipid}$ | *APM* | *t* | *fw* | $\boldsymbol{f_{FP1}}$ | $f_{lipid}$ | *APM* | *t* | *fw* | $\boldsymbol{f_{FP2}}$ | $f_{lipid}$ | *APM* | *t* | *fw* | $\boldsymbol{f_{FP4}}$ | $f_{lipid}$ | *APM* |
| Tails | 16±1 | 0 | 100 | 98±6 | 15±1 | 0 | **7** | 93 | 108±7 | 13±1 | 0 | **0** | 100 | 116±9 | 16±1 | 0 | **0** | 100 | 98±6 |
| Heads | 8 | 63±1 | 37 | 98±2 | 8 | 60±1 | **6** | 34 | 108±2 | 8 | 52±1 | **17** | 31 | 117±2 | 8 | 37±1 | **25** | 38 | 97±3 |

**Table S5**. Summary of the results obtained from the NR fittings on Langmuir lipid monolayers.

| | POPC:POPS:chol | | | | POPC:POPS:chol + **FP4** | | | | |
|---|---|---|---|---|---|---|---|---|---|
| | *t (Å)* | *fw (%)* | *$f_{lipid}$ (%)* | *APM (Å$^{-2}$)* | *t (Å)* | *fw (%)* | *$f_{FP4}$ (%)* | *$f_{lipid}$ (%)* | *APM (Å$^{-2}$)* |
| Water | 6 | 100 | 0 | / | 6 | 100 | 0 | 0 | / |
| Heads IN | 7 | 31 | 69 | 54 | 7 | 29 | 0 | 71 | 53 |
| Tails IN | 32.5 | 1 | 99 | 54 | 17 | 4 | 0 | 96 | 53 |
| Tails OUT | | | | | 15 | 6 | 4 | 90 | 63 |
| Heads OUT | 7 | 31 | 69 | 54 | 7 | 38±1.5 | 3 | 59 | 63 |

| | POPC:POPS:chol | | | | POPC:POPS:chol + **FP1** | | | | | POPC:POPS:chol + 2mM $Ca^{2+}$ | | | | POPC:POPS:chol + 2mM $Ca^{2+}$ + **FP1** | | | | |
|---|---|---|---|---|---|---|---|---|---|---|---|---|---|---|---|---|---|---|
| | *t (Å)* | *fw (%)* | *$f_{lipid}$ (%)* | *APM (Å$^2$)* | *t (Å)* | *fw (%)* | ***$f_{FP1}$*** *(%)* | *$f_{lipid}$ (%)* | *APM (Å$^2$)* | *t (Å)* | *fw (%)* | *$f_{lipid}$ (%)* | *APM (Å$^2$)* | *t (Å)* | *fw (%)* | ***$f_{FP1}$*** *(%)* | *$f_{lipid}$ (%)* | *APM (Å$^2$)* |
| Water | 5±1 | 100 | 0 | / | 5±1 | 100 | **0** | 0 | / | 4±1 | 100 | 0 | / | 5±1 | 100 | **0** | 0 | / |
| Heads IN | 7 | 29±1 | 71 | 53±2 | 7 | 34±1 | **0** | 66 | 56±2 | 7 | 34±2 | 66 | 56±3 | 7 | 32±1 | 3 | 65 | 57±2 |
| Tails IN | 33±2 | 2±0.2 | 98 | 53±9 | 17±1 | 10±0.4 | **0** | 90 | 56±6 | 17±1 | 11±0.4 | 89 | 56±5 | 17±1 | 7±0.3 | 4 | 89 | 56±6 |
| Tails OUT | | | | | 17±1 | 2±0.3 | **4** | 94 | 55±11 | 16±1 | 3±0.3 | 97 | 57±9 | 16±1 | 1±0.3 | 7 | 92 | 57±24 |
| Heads OUT | 7 | 29±1 | 71 | 53±2 | 7 | 33±2 | **0** | 67 | 56±3 | 7 | 35±2 | 65 | 57±3 | 7 | 22±1 | 12 | 66 | 56±3 |

| | POPC:POPS:chol + 10mM $Ca^{2+}$ | | | | POPC:POPS:chol + 10mM $Ca^{2+}$ + **FP1** | | | | | POPC:POPS:chol + **FP1** + EDTA | | | | |
|---|---|---|---|---|---|---|---|---|---|---|---|---|---|---|
| | *t (Å)* | *fw (%)* | *$f_{lipid}$ (%)* | *APM (Å$^2$)* | *t (Å)* | *fw (%)* | ***$f_{FP1}$*** *(%)* | *$f_{lipid}$ (%)* | *APM (Å$^2$)* | *t (Å)* | *fw (%)* | ***$f_{FP1}$*** *(%)* | *$f_{lipid}$ (%)* | *APM (Å$^2$)* |
| Water | 3±1 | 100 | 0 | / | 3±1 | 100 | **0** | 0 | / | 3±1 | 100 | **0** | 0 | / |
| Heads IN | 7 | 34±3 | 66 | 56±5 | 7 | 43±1 | **3** | 54 | 69±2 | 7 | 52±1 | **0** | 48 | 78±2 |
| Tails IN | 33±2 | 8±0.4 | 92 | 57±6 | 34±2 | 18±0.2 | **8** | 74 | 69±5 | 17±1 | 34±0.3 | **1** | 65 | 78±5 |
| Tails OUT | | | | | | | | | | 17±1 | 30±0.2 | **7** | 63 | 81±5 |
| Heads OUT | 7 | 34±3 | 66 | 56±5 | 7 | 43±1 | **3** | 54 | 69±2 | 7 | 50±1 | **4** | 46 | 80±2 |

| | POPC:POPS:chol | | | | POPC:POPS:chol + **FP2** | | | | | POPC:POPS:chol + 2mM $Ca^{2+}$ | | | | POPC:POPS:chol + 2mM $Ca^{2+}$ + **FP2** | | | | |
|---|---|---|---|---|---|---|---|---|---|---|---|---|---|---|---|---|---|---|
| | *t (Å)* | *fw (%)* | *$f_{lipid}$ (%)* | *APM (Å$^2$)* | *t (Å)* | *fw (%)* | ***$f_{FP2}$*** *(%)* | *$f_{lipid}$ (%)* | *APM (Å$^2$)* | *t (Å)* | *fw (%)* | *$f_{lipid}$ (%)* | *APM (Å$^2$)* | *t (Å)* | *fw (%)* | ***$f_{FP2}$*** *(%)* | *$f_{lipid}$ (%)* | *APM (Å$^2$)* |
| Water | 5±1 | 100 | 0 | / | 5±1 | 100 | **0** | 0 | / | 5±1 | 100 | 0 | / | 5±1 | 100 | **0** | 0 | / |
| Heads IN | 7 | 30±1 | 70 | 53±2 | 7 | 30±1 | **0** | 70 | 53±2 | 7 | 37±2 | 63 | 59±3 | 7 | 33±2 | **0** | 67 | 56±7 |
| Tails IN | 33±2 | 3±0.2 | 97 | 54±7 | 17±1 | 3±0.2 | **0** | 97 | 54±7 | 17±1 | 15±0.3 | 85 | 59±5 | 17±1 | 9±0.4 | **0** | 91 | 56±6 |
| Tails OUT | | | | | 16±1 | 9±0.2 | **0** | 91 | 60±5 | 16±1 | 1±0.3 | 99 | 53±19 | 17±1 | 1±0.4 | **0** | 99 | 53±24 |
| Heads OUT | 7 | 30±1 | 70 | 53±2 | 7 | 35±1 | **3** | 62 | 60±2 | 7 | 30±2 | 70 | 53±4 | 7 | 6±2 | **24** | 70 | 53±18 |

**Table S6**. Summary of the results obtained from the NR fittings on solid-supported bilayers

### SUPPLEMENTARY FIGURES

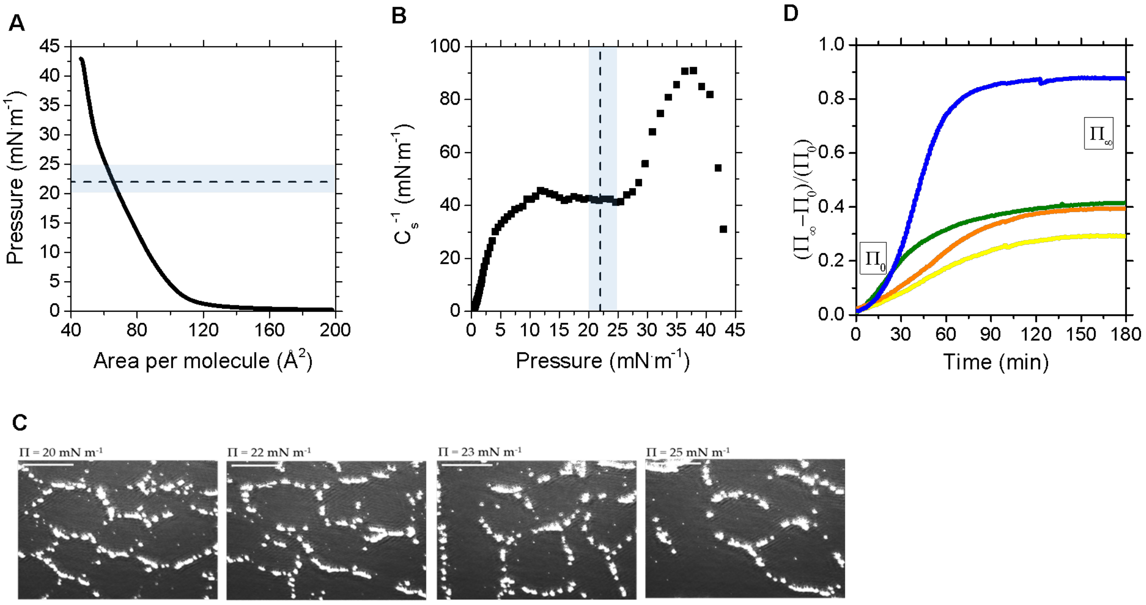


**Figure S1. A Surface Pressure isotherm for a biomimetic PM monolayer**. Surface pressure (Π) – Area isotherm of hydrogenous Plasma Membrane mimicking monolayer (left) and related compressibility modulus **B** calculated as -A(dΠ/dA). The dashed line indicates the pressure 22mN.m-1 and the shadow region the area for which the BAM images were taken. **C** Brewster Angle Microscopy (BAM) images of PM monolayer at different lateral pressures. Bright spots represent cholesterol domains. Scale bars are 50μm. **D** Profile of normalized surface pressure upon FPs interaction with PM monolayers.

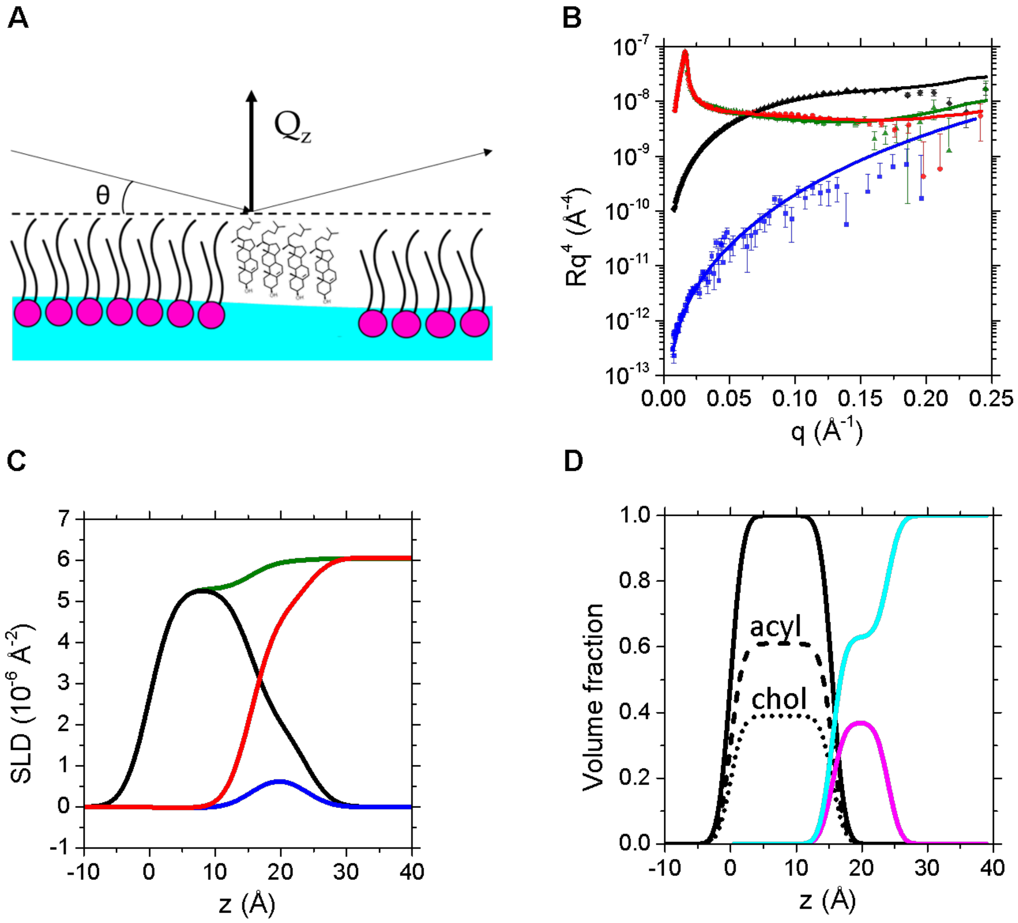


**Figure S2. Specular Neutron Reflectometry on Langmuir monolayers. A** Sketch representing a PM monolayer. Incoming and reflected neutron beams are shown. **B** Neutron reflectivity data of fully deuterated PM lipids in $D_2O$ (red diamonds) and ACMW (green circles) together with hydrogenous PM lipids in $D_2O$ (black squares) and ACMW (blue triangles). The fitting curves of deuterated PM lipids in $D_2O$ (red curve) and ACMW (green), hydrogenous PM lipids in $D_2O$ (black) and ACMW (blue) are shown. The different panels are displayed on a $Rq^4$ scale to show the agreement between the experimental and theoretical data at larger q-values. **C** Scattering length density, SLD, profiles corresponding to fits are plotted in **B**. **D** Neutron volume fraction profiles normal to the interface of monolayers to highlight the distribution of tails (black) and heads (magenta).

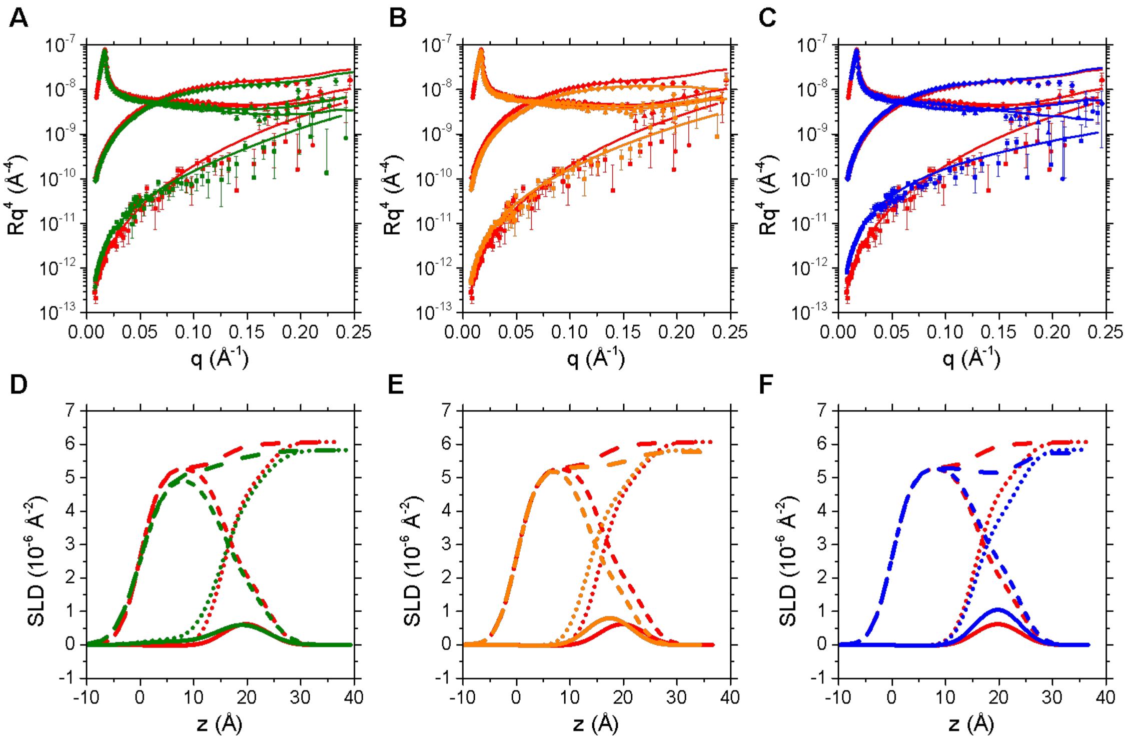


**Figure S3**. Experimental (symbols) and simulated (lines) neutron reflectivity profiles of PM monolayers, at the surface pressure value of 23 $mN.m^{-1}$, in the absence (red) and presence of **A** FP1, **B** FP2 and **C** FP4 (green, orange and blue, respectively). Data at two isotropic contrasts ($D_2O$ and ACMW (8% $D_2O$)) have been measured for hydrogenated lipid monolayers and in ACMW for deuterated version of lipid mixtures. Solid lines in A, B and C are simulated curves calculated according to a 2-layers model and the parameter listed in Table S3. Figures are displayed on a $Rq^4$ scale to show the quality of the fits at high q values. Scattering length profiles corresponding to fits of the isotropic contrasts are plotted in **D**, **E** and **F**, respectively. Continuous, long, short and dot-dot dashed lines indicate the bilayer SLD profile in ACMW and $D_2O$ isotopic contrasts with hydrogenated lipids, and in ACMW and $D_2O$ isotopic contrast with deuterated lipids, respectively.

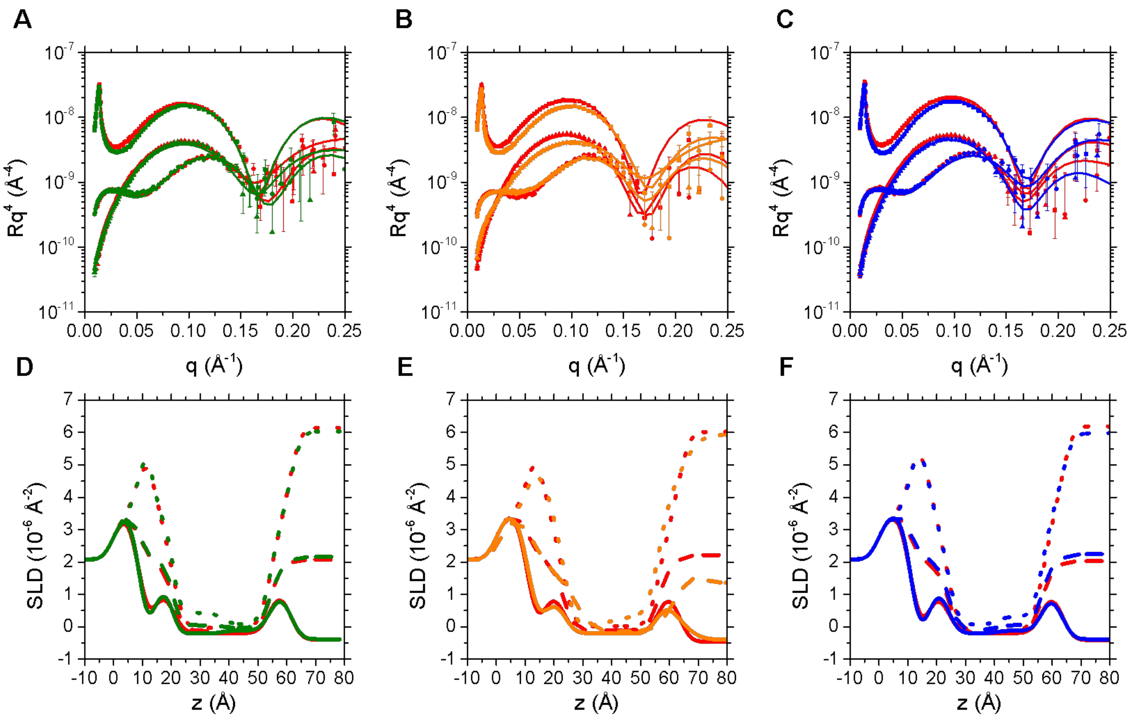


**Figure S4.** Neutron reflectivity data of PM bilayers in the absence and presence of different FP. The fitting curves in $D_2O$, ACMW, and other $D_2O$ buffers are shown. Experimental (symbols) and simulated (lines) neutron reflectivity profiles of PM solid-supported bilayers in the absence (red) and presence of FP1 (**A**), FP2 (**B**) and FP4(**C**) (green, orange and blue respectively). Data at three isotropic contrasts ($D_2O$, $H_2O$ and SiCM (38% $D_2O$)) have been measured for hydrogenated lipid bilayers. Solid lines in A, B and C are simulated curves calculated according to a 5-layers model and the parameters listed in Table S4. Figures are displayed on an $Rq^4$ scale to show the quality of the fits at high q values. Scattering length profiles corresponding to fits of the isotropic contrasts are plotted in **D**, **E** and **F**, respectively. Continuous, long, short dashed lines indicate the bilayer SLD profile in $H_2O$, SiMW and $D_2O$ isotopic contrast, respectively.

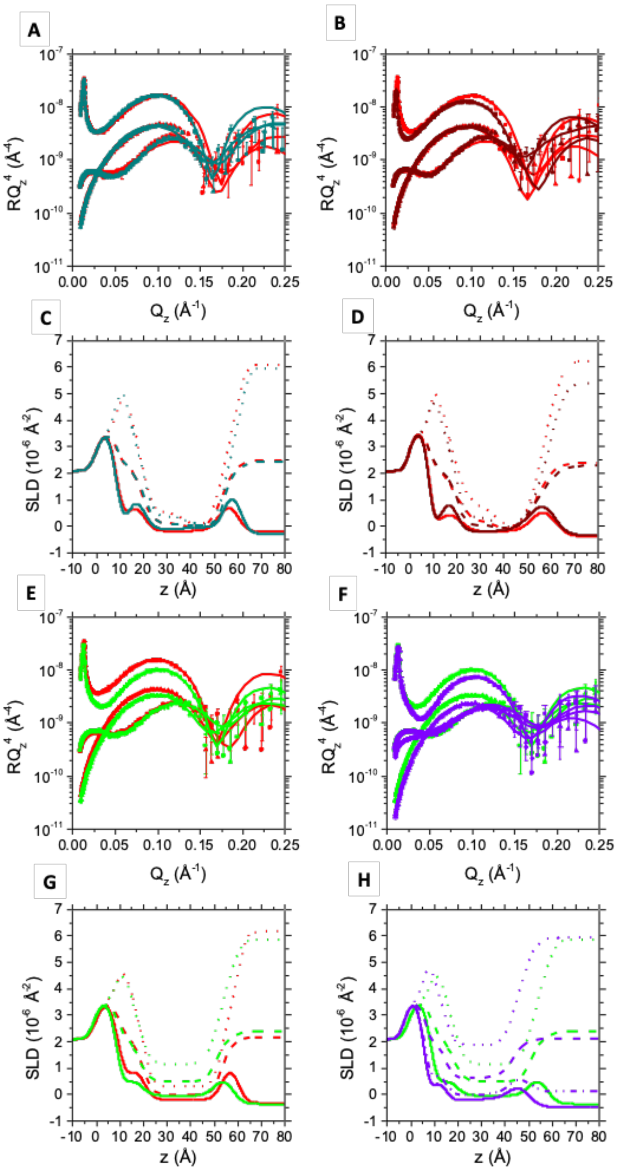


**Figure S5.** Neutron reflectivity data and resultant scattering length profiles of PM bilayers upon interaction with FP1 and FP2 in the presence and absence of $Ca^{2+}$. The fitting curves in $H_2O$, $D_2O$ and SiMW (38% $D_2O$) are shown. Experimental (symbols) and simulated (lines) neutron reflectivity and scattering length profiles of PM solid-supported bilayers in the absence (red) and presence of **A, C** FP1 in 2mM $Ca^{2+}$ (dark cyan) **B, D** FP2 in 2mM $Ca^{2+}$ (maroon), **E, G** FP1 in 10mM $Ca^{2+}$ (light green) and **F**, **H** FP1 after over-night incubation with EDTA (purple). To better characterize the system after EDTA incubation, another isotopic contrast was measured (ACMW, 8% $D_2O$). Simulated curves were calculated according to a 5-layers model and the parameters are listed in Table S4. Neutron reflectivity profiles are displayed on an $Rq^4$ scale to show the quality of the fits at high q values. Continuous, long, short and dot-dot dashed lines indicate the bilayer SLD profile in $H_2O$, SiMW, $D_2O$ and ACMW isotopic contrast, respectively.

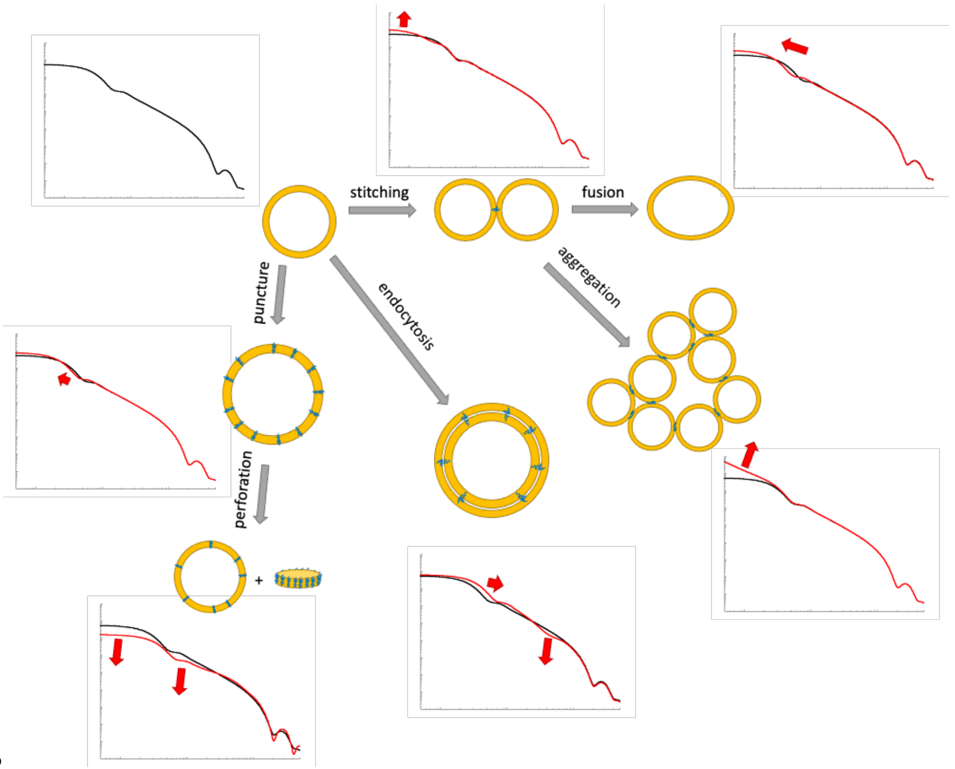
stitching
fusion
aggregation
endocytosis
puncture
perforation
+

**B**

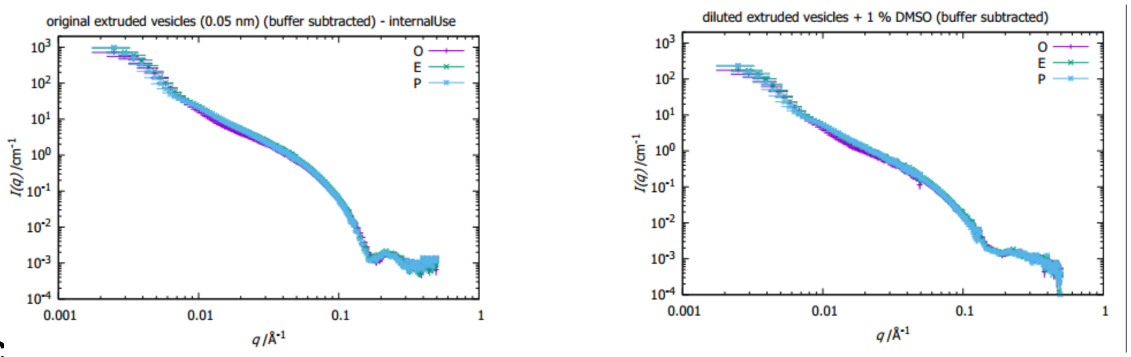
original extruded vesicles (0.05 nm) (buffer subtracted) - internalUse
diluted extruded vesicles + 1 % DMSO (buffer subtracted)
O
E
P
I(q) /cm-1
q /Å-1
0.001
0.01
0.1
1

**C**

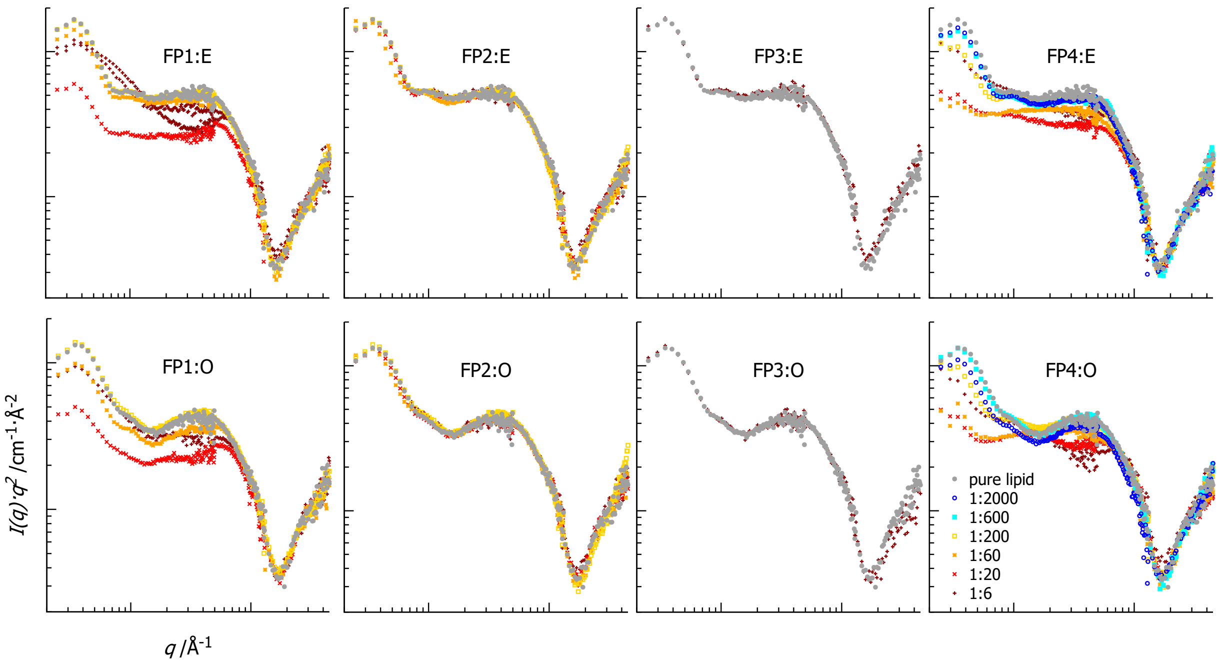
FP1:E
FP2:E
FP3:E
FP4:E
FP1:O
FP2:O
FP3:O
FP4:O
pure lipid
1:2000
1:600
1:200
1:60
1:20
1:6
I(q)·q2 /cm-1·Å-2
q /Å-1

**Figure S6. A** Sketch of observed changes in SANS scattering profiles and their interpretation. **B** Effect of 1% DMSO on diluted vesicles. While dampening the form factor contribution at high q, DMSO is not incorporated into the vesicles. **C** SANS data (D22, ILL) for extruded vesicles of "O" lipids (lacking cholesterol) and with ERGIC ("E") lipids (20% cholesterol), in Kratky plot ($I(q)q^2$ vs q). The caption indicates the mole ratio lipid:peptide. Experimentally, mixtures were prepared starting from the highest concentration of peptides, and more diluted compositions were only explored as long as effects were seen in the SANS data, which is why for FP3 only the highest ratio is available (no effect in the investigated SANS q-range, least effective peptide), while for FP4 effects were visible down to low concentrations of peptide (most effective peptide). Overall effects are relatively small by SANS (preservation of membrane content) compared to macroscopic observation (large aggregates formation). FP3 and to a lesser extent FP2 have almost no effect. FP1 and mostly FP4 show the largest effects. For FP1, the change is in line with disk formation (constant amount of membrane but decreasing amount of large vesicles), suggesting membrane perforation and at the largest peptide concentration stabilization of disks. For FP4, the increase of size observed at low q suggests binding and fusion between vesicles.

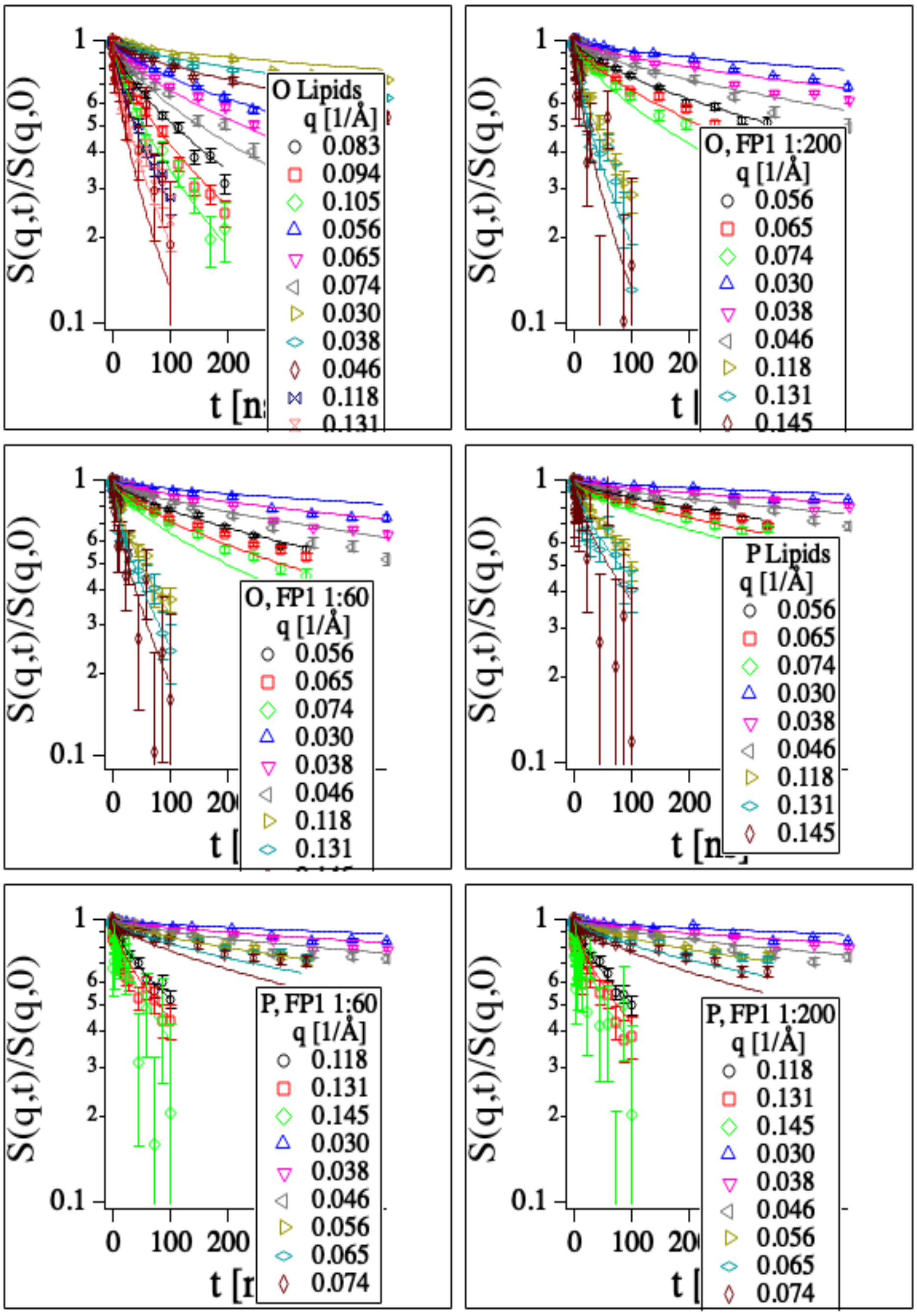


**Figure S7**. Intermediate scattering functions from NSE for samples indicated in the graphs. Good fit results are obtained with a single free parameter Γ for each sample.

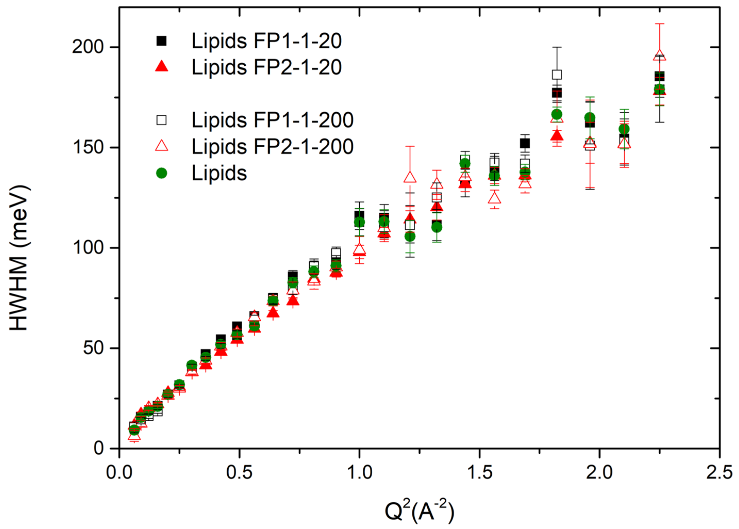


**Figure S8**. QENS-derived Lorentzian widths associated with the ensemble of peptides and lipids from the fits of data collected at 5Å on IN5 at ILL.
